# Design Guidelines For Sequestration Feedback Networks

**DOI:** 10.1101/455493

**Authors:** Ania-Ariadna Baetica, Yoke Peng Leong, Noah Olsman, Richard M. Murray

## Abstract

Integral control is commonly used in mechanical and electrical systems to ensure perfect adaptation. A proposed design of integral control for synthetic biological systems employs the sequestration of two biochemical controller species. The unbound amount of controller species captures the integral of the error between the current and the desired state of the system. However, implementing integral control inside bacterial cells using sequestration feedback has been challenging due to the controller molecules being degraded and diluted. Furthermore, integral control can only be achieved under stability conditions that not all sequestration feedback networks fulfill. In this work, we give guidelines for ensuring stability and good performance (small steady-state error) in sequestration feedback networks. Our guidelines provide simple tuning options to obtain a flexible and practical biological implementation of sequestration feedback control. Using tools and metrics from control theory, we pave the path for the systematic design of synthetic biological circuits.

## 1 Introduction

The field of synthetic biology has focused on the development of novel organisms, devices and systems for the purposes of improving industrial processes [Savage et al., 2008, Clomburg and Gonzalez, 2010, Dunlop et al., 2010, Hemme et al., 2010, Narcross et al., 2016], discovering the principles of natural biological systems [Gardner et al., 2000, Elowitz and Leibler, 2000, Guo et al., 2014], and performing biomolecular computation [Qian et al., 2011, Thubagere Jagadeesh, 2017]. Current applications of synthetic biology include industrial fermentation [Georgianna and Mayfield, 2012], toxin detection [McBride et al., 2003, Wang et al., 2009], biosensor development [Hsiao et al., 2016], diagnostics detection [Pardee et al., 2014], materials production [Cherny and Gazit, 2008, Moon et al., 2010, DeLoache et al., 2015], novel protein design [Dahiyat and Mayo, 1996, Arnold, 1998, Kuhlman et al., 2003], and biological computation [Moon et al., 2012, Daniel et al., 2013]. The applications and capabilities of synthetic biology are still expanding because it is a relatively novel field. However, synthetic systems have a unique set of challenges and limitations.

An important goal of synthetic biology is to engineer reliable, robust, and well-performing systems from standardized biological parts that can easily be combined together [Canton et al., 2008, Arkin, 2008, Kwok, 2010, Khalil and Collins, 2010, Qian and Del Vecchio, 2018]. Nonetheless, synthetic systems can lack robustness and be sensitive to their biological implementation [Yeung et al., 2014, Del Vecchio, 2015, Yeung et al., 2017]. Differences in their biological parts, different model organism implementations [Klumpp et al., 2009], or different experimental conditions can cause synthetic systems to cease to function properly [Purnick and Weiss, 2009, Gomez et al., 2016]. This limits their applicability, as well as the engineering of more complex synthetic systems.

The development of synthetic biological systems is limited by the lack of consistent functionality, performance, and robustness. These limitations are also present in mechanical and electrical engineering. However, tools and concepts have been developed to ameliorate them [Levine, 2010, Friedland, 2012, Doyle et al., 2013, Aström and Murray, 2008]. For example, the engineering design cycle described in Figure 1 is a framework for the iterative design, build, testing, and learning of engineered systems. The engineering design cycle can also be iteratively applied to synthetic systems to achieve the desired performance standards. In this paper, we employ engineering principles and tools to the design of a class of well-performing, robust synthetic biological systems.

**Figure 1:**
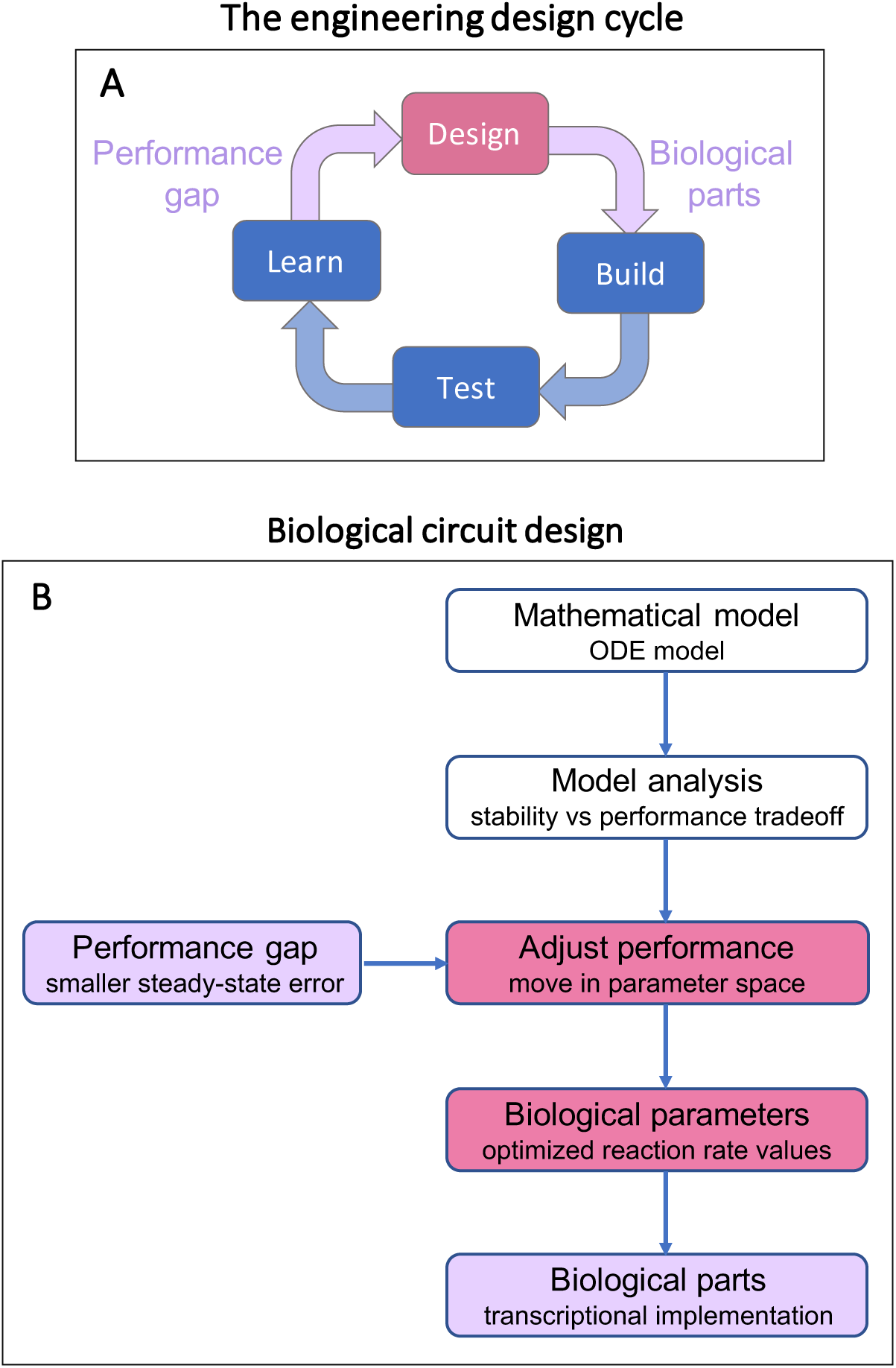
The engineering design cycle is an iterative approach to improving the performance of engineered and biological systems. **A.** The engineering design cycle is an approach by which an engineering or biological system is iteratively designed, built, tested, and learned until it becomes sufficiently improved in performance. When there is a gap in performance, we find a new biological design and associated biological parts for its implementation. **B.** An example of a performance gap is when the biological system’s steady-state error is larger than desired. Provided that the structure of the biological system is fixed, this indicates a necessary adjustment in the system’s performance by moving across its parameter space. Thus, we find optimized reaction rate values that result in the desired performance specifications of small steady-state error. Using the optimized reaction rate values, we can determine corresponding biological parts that provide this particular implementation. In this paper, we design the sequestration feedback network in Figure 2 to fulfill stability and performance specifications. In turn, these performance specifications inform the biological implementation of sequestration feedback networks. Throughout this paper, we use the stability and performance properties of sequestration feedback networks determined in [Baetica, 2018, Olsman et al., 2018, Briat et al., 2016] to inform their design.

**Figure 2:**
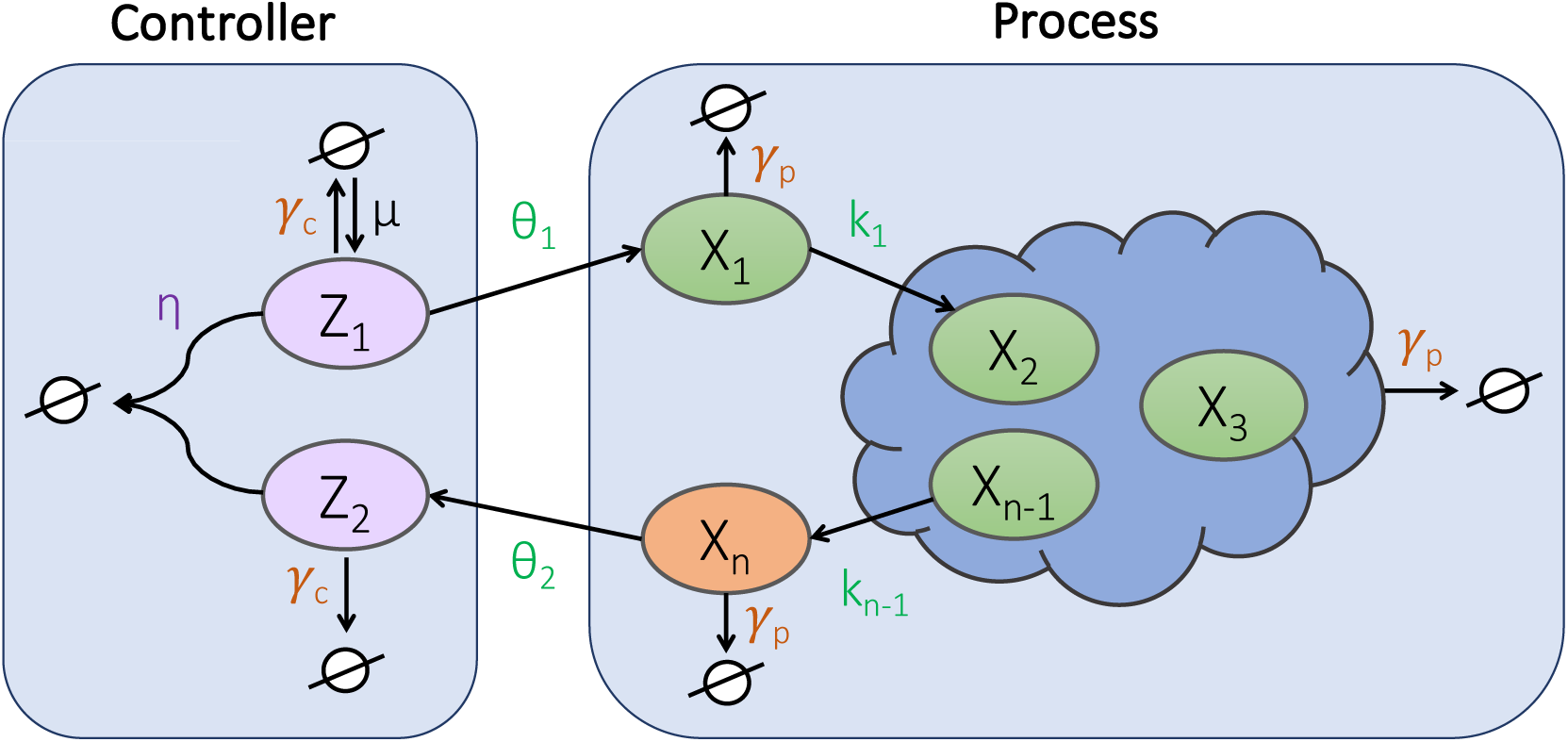
The sequestration feedback network’s controller and process networks. The goal of the sequestration controller is for the process output species *X_n_* (orange) to track the reference signal. The reference acts on the sequestration feedback network through the constitutive production of controller species *Z*_1_ (purple) at rate *µ*. The controller species *Z*_1_ and *Z*_2_ (purple) bind together at rate *η* in a sequestration reaction to form an inactive complex represented by the empty set. The controller species *Z*_1_ acts on the process input species *X*_1_ at rate *θ*_1_. The process output species *X_n_* acts on the controller species *Z*_2_ at rate *θ*_2_. Process input species *X*_1_ creates process species *X*_2_ at rate *k*_1_ and process species *X*_*n* − 1_ creates process output species *X_n_* at rate *k*_*n* − 1_. Additionally, process species *X*_2_ …, *X*_*n* − 1_ (green) also interact with each other in unimolecular or bimolecular reactions. The controller and the process species are subjected to degradation and dilution, which is indicated by arrows pointing to empty sets. Rates *γ_c_* and *γ_p_* each encompass both degradation and dilution. This diagram extends the setup in [Briat et al., 2016] by the inclusion of the controller species’ degradation and dilution.

A widely used tool to improve the performance of mechanical and electrical systems is feedback control. Feedback control allows a system to take corrective action based on the measured differences between the current and desired performance [Levine, 2010, Aström and Murray, 2008]. The foremost benefit feedback control provides to biological, mechanical, and electrical systems is robustness to uncertainty. Should the system undergo a change such an external disturbance, the feedback controller ensures that the system retains good performance properties such as small steady-state error and fast response time by correcting for the change. Additionally, feedback control stabilizes an unstable process and accelerates a slow process. Yet, if poorly designed, feedback control can inadvertently amplify noise and exacerbate instability [Levine, 2010, Del Vecchio and Murray, 2015, Aström and Murray, 2008].

Feedback is ubiquitous in natural biological systems, where it serves to regulate their behavior. Examples of feedback control found in natural biological systems include the regulation of body temperature [Werner, 2010], circadian rhythms [Rust et al., 2007], calcium [El-Samad et al., 2002], and glycolysis [Chandra et al., 2011]. In this paper, we explore the use of a feedback controller for synthetic biological systems that relies on the sequestration reaction of two biological species. This sequestration feedback controller is illustrated in Figure 2. Examples of synthetic systems that use sequestration feedback include the concentration tracker in [Hsiao et al., 2014], the two bacterial growth controllers in [McCardell et al., 2017], and the gene expression controller in [Annunziata et al., 2017]. An example of a natural sequestration feedback system uses sigma factor *σ*^70^ and anti-sigma factor Rsd [Jishage and Ishihama, 1999] for the two sequestering species.

The sequestration feedback introduced in Figure 2 has been a promising implementation of feedback control that can achieve perfect adaptation. Perfect adaptation (whose engineering counterpart is integral control) ensures adaption to disturbances with no steady-state error. A biological system with perfect adaptation displays excellent robustness and performance in terms of steady-state error. Perfect adaptation is a desirable property that can be found both in natural biological systems and in synthetic systems [Ferrell Jr, 2016, Goentoro et al., 2009, El-Samad et al., 2002, Yi et al., 2000, Del Vecchio and Murray, 2015]. Briat *et al.* [Briat et al., 2016] studied stochastic sequestration feedback systems with no controller species degradation and demonstrated that they can achieve perfect adaptation under certain conditions of stability. Nevertheless, we and others have found the assumption of zero controller species degradation to be too restrictive for the practical implementation of sequestration feedback [Ang et al., 2010, Ren et al., 2017, Qian and Del Vecchio, 2018, Olsman et al., 2018].

Hence, current synthetic sequestration controllers may or may not be able to achieve perfect adaptation in a practical biological implementation. In Section 2.2.2, we suggest another possible implementation of sequestration feedback that accounts for the controller species’ degradation and dilution and ensures zero steady-state error if the closed loop system is stable. Nevertheless, this implementation depends on an exact relationship between the controller and the process network parameters, which renders it also inflexible and impractical. In Section 2.3, we relax the requirement that the sequestration controller achieve perfect adaptation and we simulate and design sequestration feedback controllers with small steady-state error, large stability margin, and good disturbance rejection properties. These controllers are advantageous because they do not require an extremely precise implementation.

Previous research on sequestration feedback networks has determined conditions for their stability, along with their performance and robustness properties [Briat et al., 2016, Qian and Del Vecchio, 2018, Baetica, 2018, Olsman et al., 2018]. The current paper uses these properties of stability, performance, and robustness to find optimal designs for sequestration feedback networks. In Section 2.1, we introduce a model of sequestration feedback networks with controller species degradation and dilution. Then we summarize and expand their properties of stability, performance, and robustness of sequestration networks in Section 2.2. In Section 2.3, we discuss how the choice of biological parts impacts these properties. Additionally, we solve a case study design problem for a sequestration feedback network with two process species. Finally, we develop general guidelines for design of sequestration controllers when the process network is already specified by challenging experimental constraints.

## 2 Results

### 2.1 Modeling Sequestration Feedback Networks with Controller Species Degradation

In this section, we introduce the sequestration feedback network’s model and chemical reactions and we clarify our assumptions about its dynamics. Additionally, we outline the properties of sequestration feedback controllers. In Figure 2, we illustrate a general sequestration feedback network with controller species degradation and dilution. The controller network consists of two biochemical species *Z*_1_ and *Z*_2_ that sequester each other into an inactive complex. This sequestration reaction computes the difference in concentration between the two controller species. If there are more molecules of *Z*_1_ than *Z*_2_, then the sequestration feedback encodes the quantity *Z*_1_ − *Z*_2_, whereas if there are more molecules of *Z*_2_ than *Z*_1_, then it encodes the quantity *Z*_2_ − *Z*_1_. The controller network is connected to the process network through its interactions with process species *X*_1_ and *X_n_*. The goal of the controller network is to ensure that the output species *X_n_* of the process network tracks a reference signal set by rates *µ* and *θ*_2_. As we explain in this section, the difference between controller species *Z*_1_ and *Z*_2_ encodes the error between process output species *X_n_* and the reference signal.

Here, we assume that the two controller species *Z*_1_ and *Z*_2_ sequester each other into an inactive complex at rate *η*. The reference signal comes in through controller species *Z*_1_’s constitutive production at rate *µ*. Controller species *Z*_1_ acts on the process network input species *X*_1_ at rate *θ*_1_, while the process network output species *X_n_* acts on the second controller species *Z*_2_ at rate *θ*_2_. Indeed, species *Z*_1_ is the actuator and species *Z*_2_ is the sensor for the process network. For simplicity, we assume that the controller species are degraded at the same rate *γ_c_* and that the process species are degraded at an equal rate *γ_p_*. Under these assumptions and notation, the chemical reactions that describe the sequestration feedback network are as follows:
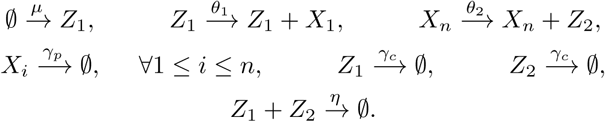

For simplicity, we omit the chemical reactions between process species *X*_2_, …, *X*_*n* − 1_ for the rest of this section. We let *t* denote time and *x*_1_, …, *x_n_* denote the concentrations of the process species *X*_1_, …, *X_n_*. We let *z*_1_ and *z*_2_ denote the concentrations of the controller species *Z*_1_ and *Z*_2_, respectively. Thus, the sequestration controller can be modeled as follows:
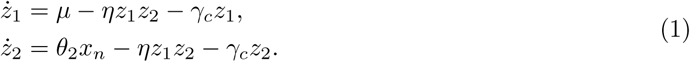

We subtract the dynamics of the two controller species and obtain the following equation:
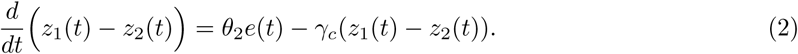

If we define the error signal as 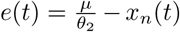, then the control action *z*_1_(*t*) − *z*_2_(*t*) integrates the error signal *e*(*t*) as follows:
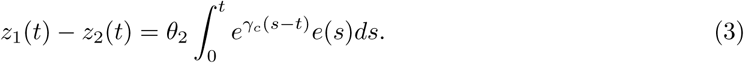

If the closed loop sequestration feedback network is stable, then equations (1) have a steady-state and we can evaluate the magnitude of the steady-state error. Moreover, when the controller species do not degrade (*γ_c_* = 0), then the sequestration feedback network exhibits the property of perfect adaptation since *µ* = *θ*_2_*x_n_* at steady-state. This implies implies that *x_n_* perfectly tracks the error signal 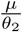. Perfect adaptation is a desirable property of biological systems because it guarantees zero steady-state error and robustness to disturbances, irrespective of the process network as long as the closed loop network remains stable. This allows for an imprecise implementation of the process network, although the zero controller degradation rate requires a very challenging implementation of the sequestration controller species.

When we assume stability of the closed loop sequestration network and a nonzero degradation rate of the controller species (*γ_c_* > 0), equation (3) demonstrates that the sequestration controller is a lag compensator and that it integrates the error signal [Aström and Murray, 2008, Ren et al., 2017]. When the controller species degradation and dilution rates are small, the sequestration controller approaches integral control, whereas when the controller species degradation and dilution rates are large, it approaches proportional control. A detailed comparison of sequestration feedback networks with and without controller species degradation and dilution is available in the supplement (Section S4.1).

In particular, it is not immediately apparent whether the sequestration controller with controller species degradation and dilution retains the property of zero steady-state error. We know that while integral control guarantees zero steady-state error under a variety of process network implementations, the performance of a lag compensator depends on the parameters of the sequestration feedback network. We analyze the performance of sequestration feedback networks with controller species degradation and dilution in Section 2.2.2.

We briefly state the stability criteria for the closed loop sequestration feedback networks with zero and nonzero controller species degradation rates in Section 2.2. An in depth discussion of these stability criteria is presented in [Baetica, 2018, Olsman et al., 2018]. We also note that the stability criteria change between sequestration feedback networks with and without controller species degradation and dilution.

Since both the process and the controller networks can only be imprecisely built with biological parts, we consider how this affects the properties of stability and performance of sequestration feedback networks in Section 2.3. In particular, we replace the stringent performance requirement of zero steady-state error with a more flexible and practical performance requirement of small steady-state error. In Section 2.3, we formulate guidelines for implementing sequestration feedback networks with large stability margin and small steady-state error.

### 2.2 Stability Criteria and Performance Goals for Sequestration Feedback Networks

Throughout the rest of the paper, we restrict the discussion to the class of sequestration feedback networks in Figure 3, unless otherwise specified. The process network is simplified from Figure 2 to only include catalytic reactions between each process species *X_i_* and process species *X_i_*_+1_ for 1 ≤ *n* − 1 and degradation reactions with equal rates *γ_p_*. No other interactions are present between process species *X*_1_, …, *X_n_*. Accordingly, we can model the sequestration feedback network in Figure 3 using the following system of equations:
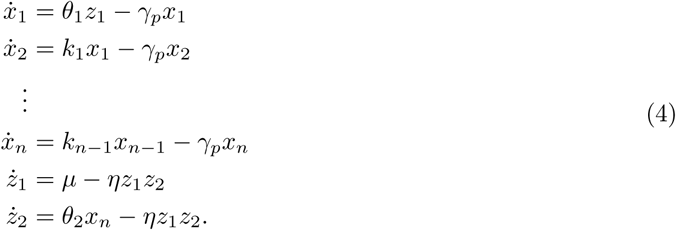

**Figure 3:**
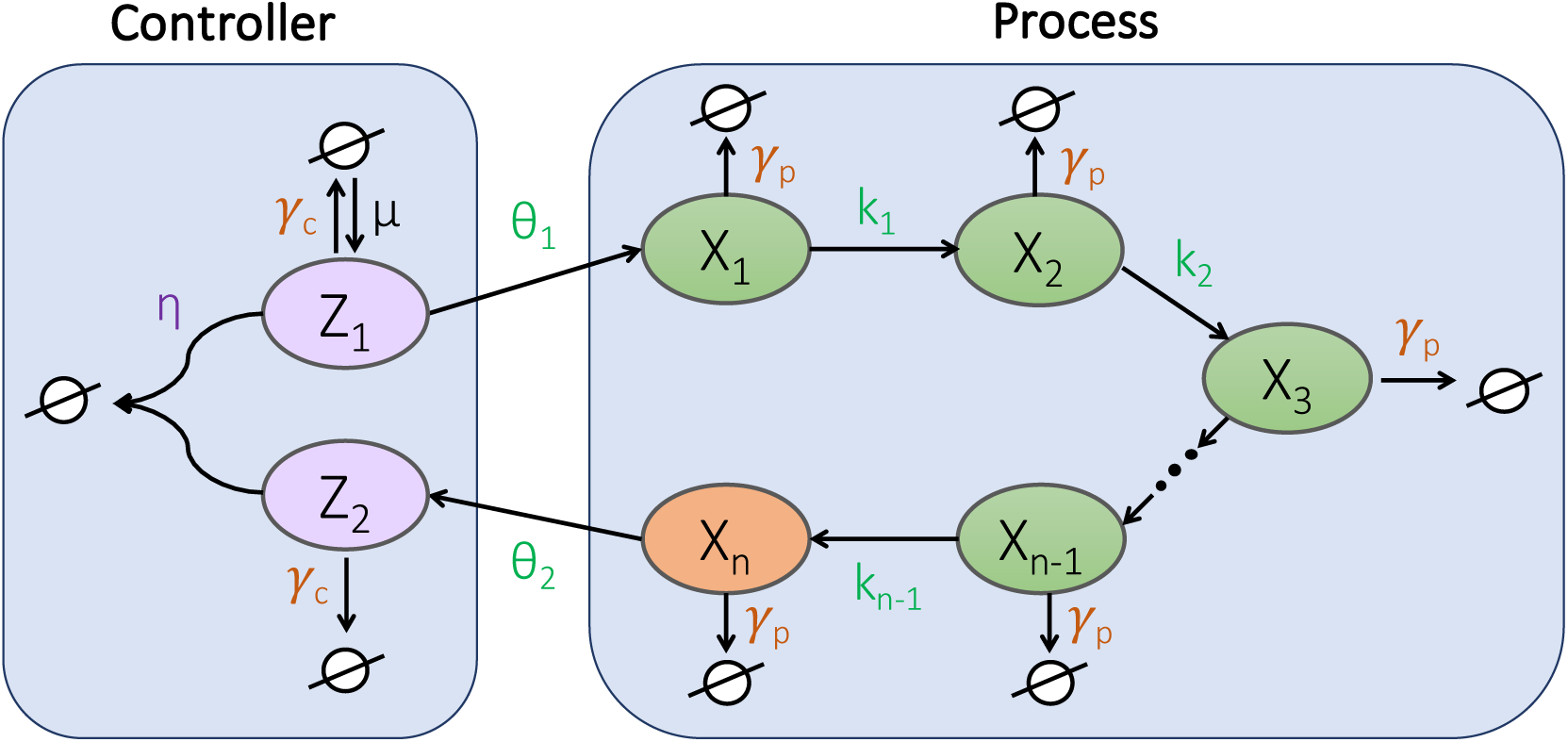
A class of sequestration feedback networks with simplified process networks. This class of sequestration feedback networks represents a subset of the networks in Figure 2. The process networks are simplified to only allow bimolecular chemical reactions between species with consecutive numbering; each process species *X_i_* is created by the previous process species *X*_*i* − 1_ for 2 ≤ *i* ≤ *n*. For simplicity, we assume that the process species degradation rates *γ_p_* (orange) are equal. Similarly, we assume that the controller species degradation rates *γ_c_* (orange) are equal. The sequestration reaction rate *η* is in purple and the process species production rates *θ*_1_, *θ*_2_, *k*_1_, … *k*_*n* − 1_ are in green. An example process network that follows these assumptions implements the process species as proteins inside a bacterial cell. These simplifying assumptions match the setup in [Baetica, 2018, Olsman et al., 2018].

This particular class of sequestration feedback networks is amenable to the derivation of an analytical stability criterion for the closed loop sequestration feedback network, as well as to the derivation of an analytical form of their steady-state error under the assumptions of “strong sequestration feed-back” [Baetica, 2018, Olsman et al., 2018]. The strong feedback assumptions ensure that we can find the complex roots of the characteristic polynomial associated with the linearized sequestration feedback network. Therefore, we can determine if the closed loop sequestration feedback network is stable based on the real parts of the roots of their characteristic polynomial. We introduce the following notation to restate the strong feedback assumptions and the stability criteria of the sequestration feedback networks in Figure 3:
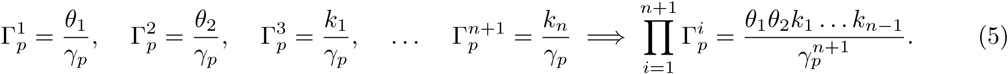

The quantities 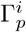, 1 ≤ *n* + 1, are all constants since the controller and process network rates *θ*_1_, *θ*_2_, *k*_1_, …, *k_n_*, and *γ_p_* have units of s^−1^. For *n* > 2, under the strong feedback assumptions, the stability conditions for the sequestration feedback network in Figure 3 are restated in Table 1 from [Baetica, 2018, Olsman et al., 2018]. Intuitively, the strong feedback assumptions ensure that the controller network can react faster than the dynamics of the process network. Deriving the analytical stability criteria when *n* > 2 relies on comparing the process and the controller species’ degradation rates. In the three cases outlined in Table 1, the sequestration feedback network is stable if and only if the quantity 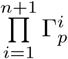 is bounded by a constant that only depends on *n*, the number of process species.

**Table 1:**
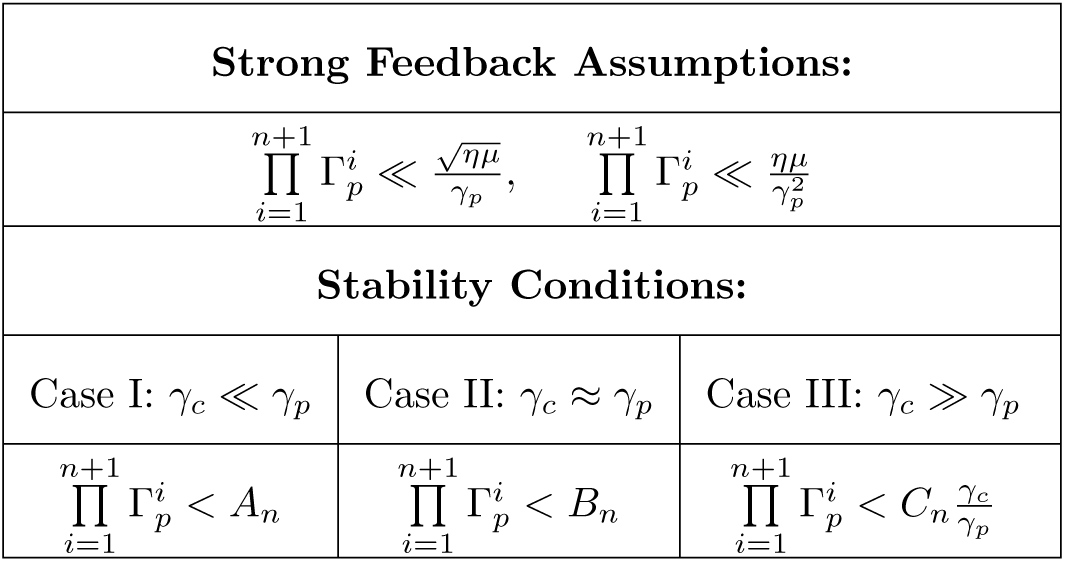
Stability Depends on the Controller and the Process Species’ Production and Degradation Rates. Under the strong feedback assumptions, there are analytical criteria for the stability of the sequestration feedback networks in Figure 3. Deriving the analytical stability criteria for a sequestration network with at least three process species relies on comparing the process and the controller species’ degradation rates. In Cases I and II, the sequestration feedback network is stable if and only if the product of the ratios of production and degradation rates is smaller than the constants *A_n_* and *B_n_*, respectively. These constants do not depend on the parameters of the sequestration network, only on the number of process species. In Case III, stability depends on the constant *C_n_*, as well as on the ratio between the controller and the process species degradation rates. These analytical stability criteria are derived in [Baetica, 2018, Olsman et al., 2018].

When the process network has only two species (*n* = 2), we can derive an analogous analytical criterion. Under the strong feedback assumptions of 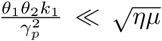 and 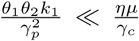, stability is achieved if and only if
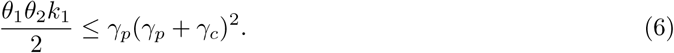

In Section 2.2.1, we extend these stability results from [Baetica, 2018, Olsman et al., 2018] to include sequestration feedback networks with process species that are degraded at different rates.

Moreover, we consider the robustness of sequestration feedback networks for *n* = 2. According to [Aström and Murray, 2008], a measure of robustness is the infinity norm of the sensitivity function, which represents the worst-case disturbance amplification for the system to an oscillatory input. Analytically computing the infinity norm of the sensitivity function can be challenging, particularly for *n* > 2. Thus, [Olsman et al., 2018] computes a lower bound 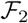 to the infinity norm of the sensitivity function as:
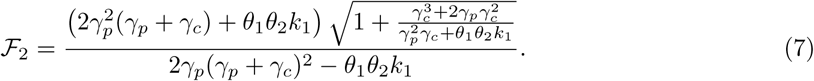

Equality in this bound is achieved when the sequestration feedback network goes from being stable to unstable. This is equivalent to attaining equality in equation (6). In Section 2.3, we refer to 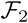 as the “fragility” of a sequestration feedback network and we use its analytical expression to design robust sequestration control.

#### 2.2.1 The Slowest Process Species Degradation Rate Impacts Stability

Here, we assess the impact of the slowest process species degradation rate on a stable sequestration feedback network. For simplicity, we consider a process network with only two species that are degraded at different rates, as illustrated in Figure 4A. We assume strong sequestration feedback and we compare the impact of two large process species degradation rates of 2 h^−1^ (panel B) to a large and a small process species degradation rate of only 0.25 h^−1^ (panel C). Clearly, the slower process species degradation rate drives the sequestration feedback network to instability.

**Figure 4:**
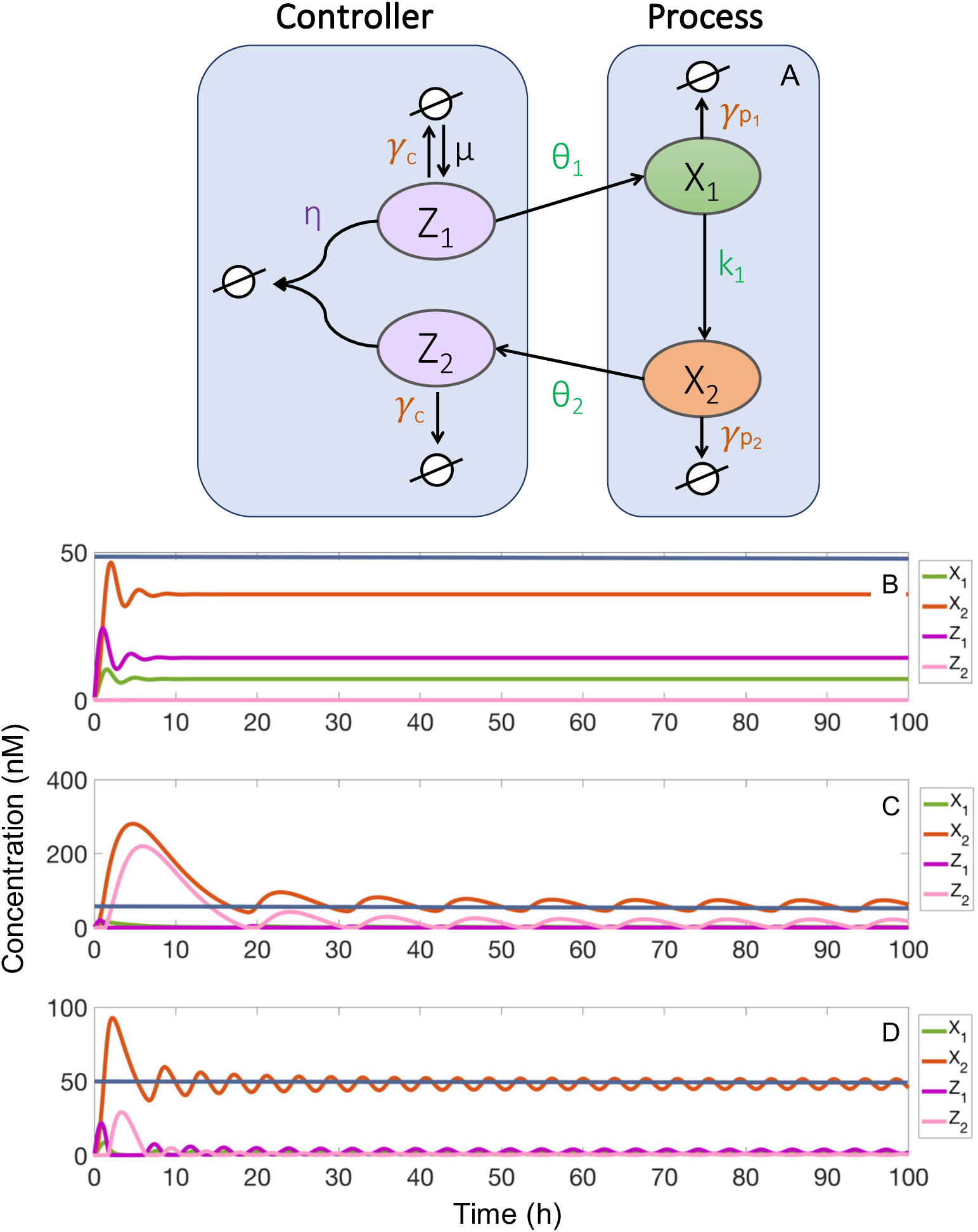
The stability of the sequestration feedback network with different process species degradation rates. **A.** We illustrate the sequestration feedback network with two different process species degradation rates, *γ_p_*_1_ and *γ_p_*_2_. Then we analyze the stability of sequestration feedback networks by assuming strong sequestration feedback, indicative of a sigma factor implementation of the controller species. **B.** We use large process species degradation rates of 2 h^−1^. We note stability since the process output species *X*_2_ (orange) tracks the reference concentration of 50 nM (dark blue), albeit with a large steady-state error. **C.** We use a process species with a small degradation rate (0.25 h^−1^) and a process species with a larger degradation rate (2 h^−1^). This results in instability, as evidenced by the sustained oscillations in the process output species *X*_2_. **D.** Both process species have small degradation rates of 0.25 h^−1^ that also result in instability. The other parameters used in these simulations are: 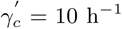, *η*′ = 50 nM^−1^, *µ*′ = 50 nM h^−1^, 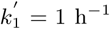, 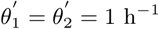.

In the supplement (Section S4.2), we derive an analytic stability criterion for the sequestration feedback network with two process species with different degradation rates. This stability criterion in presented in Theorem S1. When the sequestration feedback network in Figure 4A satisfies the strong feedback assumptions, it is stable in closed loop if and only if
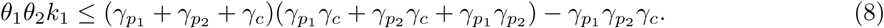

This result advances our understanding of the stability of sequestration feedback networks. We have not yet derived an analytical stability criterion for sequestration feedback networks with *n* process species with different degradation rates for *n* > 2. In the rest of the paper, we describe the performance properties of sequestration feedback networks and highlight the challenge between achieving both good stability and performance properties.

#### 2.2.2 Strong Sequestration Feedback Results in Nonzero Steady-State Error

In Section 2.1, we indicated that sequestration feedback networks with nonzero controller species degradation rates can have a nonzero zero steady-state error. Indeed, depending on the parameters of the sequestration feedback network, its steady-state error can either monotonically increase with the controller species degradation rate or it can equal zero for a specific “critical” controller species degradation rate. In particular, under the strong feedback assumptions, the steady-state error will increase monotonically with the controller species degradation rate. Therefore, an implementation of sequestration feedback with controller species degradation and dilution that satisfies the strong feedback assumptions will inevitably result in a nonzero steady-state error. Thus, the performance objective of zero steady-state error is a stringent and potentially impractical requirement. When we perform the design of sequestration feedback networks in Section 2.3.1, we relax our performance objective from a zero steady-state error to a small steady-state error. This affords us numerous options for the practical implementation of sequestration feedback networks.

For the sequestration feedback networks in Figure 2, we give conditions for the existence of a “critical” controller species degradation rate that ensures zero steady-state error in the supplement (Section S4.3). For the particular class of sequestration feedback networks in Figure 3, the critical controller species degradation rate is given as:
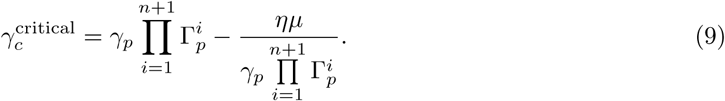

For this class of sequestration feedback networks, the critical controller species degradation rate 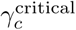 exists if and only if the process species degradation rate satisfies the inequality
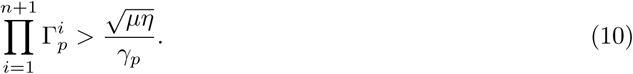

Clearly, this inequality contradicts the first strong feedback assumption in Table 1. According to [Olsman et al., 2018], when the strong feedback assumptions hold, the steady-state error is simply a monotonic function of the controller species degradation rate as follows:
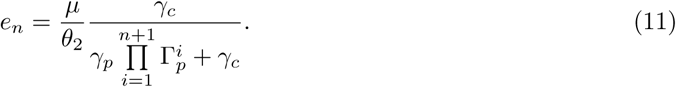

Thus, under the strong feedback assumptions, the steady-state error is a monotonically increasing function of the controller species degradation rate *γ_c_* and a monotonically decreasing function of the process species degradation rate *γ_p_*. We further discuss and illustrate the stability margin and the relative steady-state error as functions of the process and the controller species degradation rates in Section 2.3.2.

### 2.3 Developing Implementation Guidelines for Sequestration Feedback Networks

In this section, we connect our mathematical understanding of sequestration feedback networks to their biological implementation. In Section 2.2, we have discussed how different controller and process species and the chemical reactions between them affect the stability and the performance of sequestration feedback networks. In particular, the controller species sequestration rate, as well as the process and controller species degradation rates are important parameters. We use the analysis in Section 2.2 to provide guidelines for the biological implementation of robust, well-performing sequestration feedback networks.

Depending on the choice of controller species, the sequestration reaction strength and process and controller species’ degradation rates can vary over several orders of magnitude. Examples of possible sequestration controller species include transcriptional parts such as an mRNA and antisense RNA pair, protein parts such a sigma and anti-sigma pair or a toxin and antitoxin pair. First, we note that the stability of sequestration feedback networks can be improved by increasing either the degradation rates of the process or of the controller species. Intuitively, this is similar to adding more dissipation to a physical system [Aström and Murray, 2008]. However, in practice, there are limits to how high the degradation rates of the process or the controller species can be and we report representative values for these rates in Figure 5. Second, we highlight how the process and the controller species’ degradation rates influence the steady-state error of the sequestration feedback network. For a particular class of sequestration feedback networks that fulfill the strong feedback assumptions, we use the analytic expression of the steady-state error in equation (11) to bound it by tuning these degradation rates.

**Figure 5:**
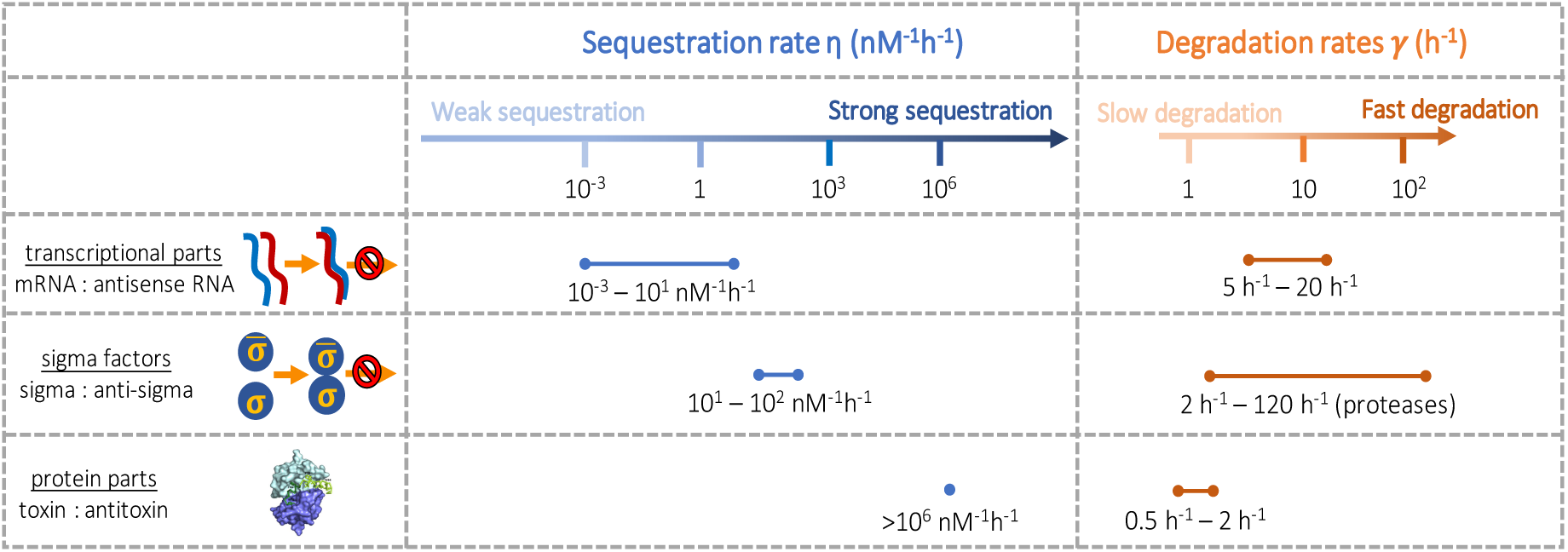
The sequestration and the degradation rates of representative biological parts that can be used in sequestration feedback networks. The sequestration reaction between the controller species can be implemented using a variety of biological parts. Example transcriptional parts are mRNA and antisense RNA. Antisense RNA inhibits the translation of complementary mRNA by base pairing to it and physically obstructing the translation machinery of the cell. Anti-sigma factors bind sigma factors to inhibit transcriptional activity. Protein parts include the toxin-antitoxin module CcdA-CcdB in *E. coli*. When CcdB outlives CcdA, it kills the cell by poisoning the DNA gyrase. The antitoxin CcdA blocks the activity of the toxin CcdB by binding together into a complex, thus allowing cells to grow normally. We include representative ranges of the sequestration reaction rate (*η*) for transcriptional parts [Walton et al., 2002], sigma factors [Sharma and Chatterji, 2008], and toxin parts [De Jonge et al., 2009]. For small *η* values, the sequestration reaction is considered weak and for large values, it is considered strong. Analogously, we give representative values of the degradation rates of the process and the controller species implemented using transcriptional parts, sigma factors, and protein parts. The degradation rate of mRNA is between 2 h^−1^ and 20 h^−1^ [Bernstein et al., 2004, Miller et al., 2011, Bernstein et al., 2002] and the synthesis rate is between 0.1 and 10 h^−1^ in yeast [Miller et al., 2011]. Toxin Hok in the type I toxin-antitoxin pair in *E. coli* is degraded at a rate of 2 h^−1^ [Steif and Meyer, 2012]. The degradation rate of the sigma factor RpoS is 1.39 h^−1^ when the *E. coli* cells are in stationary phase at 37°*C* or under stress conditions [Zhou and Gottesman, 1998]. In the presence of proteases ClpXP, this rate increases up to 120 h^−1^. The antitoxin CcdA is degraded in wild-type cells with a rate of 1.39 h^−1^ in the absence of toxin CcdB and a rate of 0.69 h^−1^ when bound in a complex with toxin CcdB [De Jonge et al., 2009]. Additional information about the representative rate constants is provided in Section S4.4.1. We also note that protein synthesis rates can be modified by varying promoters and ribosome binding sites. This figure includes a partial reproduction of a figure in [De Jonge et al., 2009] with permission from the author.

Using our analytical results for stability and steady-state error, we provide general guidelines for the implementation of sequestration feedback networks. We suggest simple tuning options to improve the stability, the robustness, and the performance of sequestration networks. When the process network is already specified by experimental constraints, we explain how to choose a sequestration controller that guarantees good stability and performance properties (Section 2.3.2).

#### 2.3.1 The Choice of Biological Parts Impacts the Performance of Sequestration Feedback Networks

Sequestration feedback networks can be built using a variety of biological parts for the two sequestering controller species. Several examples of parts for the two controller species are illustrated in Figure 5. They include transcriptional parts such a mRNA and antisense RNA pair, sigma factors such a sigma and anti-sigma pair, or protein parts such as a toxin and antitoxin pair. Transcriptional parts can be obtained from systems such as the hok-sok type I toxin-antitoxin system in *E. coli* or from parts already mined for synthetic biological systems. The hok gene product is a toxin that kills cells without its antidote, the antisense RNA sok that is complementary to the hok mRNA [Gerdes, 1988]. A synthetic system of RNA and antisense RNA that originally performed translation initiation control is adapted to regulate transcriptional elongation in [Liu et al., 2012]. Protein parts can be obtained from type II toxin-antitoxin bacterial systems such as CcdA-CcdB. The toxin CcdB targets DNA gyrase and induces the breaking of DNA and subsequently cell death, while its antitoxin CcdA inhibits CcdB toxicity by sequestering it into a very stable CcdA-CcdB complex [De Jonge et al., 2009]. Protein parts can also be sigma and anti-sigma factors such as *σ*70 binding to Rsd and inhibiting RNA polymerase, which slows transcription and inhibits *E. coli* growth [Sharma and Chatterji, 2008].

Depending on the choice of parts for the controller species in sequestration feedback networks, the binding affinity of the sequestration reaction can vary over several orders of magnitude, as illustrated in Figure 5. Yet, the binding affinity of the sequestration reaction influences the stability of the sequestration feedback network, as discussed throughout Section 2.2. Accordingly, we need to carefully consider whether our implementation choice for the controller parts results in stable closed loop control.

Additionally, depending on the choice of implementation of the process and the controller species, their degradation rates can also vary, as illustrated in Figure 5. Typically, the half-life of proteins inside the cell is long since they are slowly degraded at a rate between 1 h^−1^ and 2 h^−1^. However, the half-life of mRNA inside the cell is very brief, so we must include this degradation rate in our model. The degradation rate of the controller species influences the stability of the sequestration feedback network, as discussed in Section 2.2.

Therefore, we refer to Figure 5 for biologically representative values for the controller species’ sequestration rate and for the process and the controller species’ degradation rates throughout Section 2.3. In the next section, we use these reaction rate values to choose biological parts for the implementation of sequestration controllers such that the sequestration feedback network has good properties of stability, robustness, and performance.

#### 2.3.2 Developing Guidelines for the Design of Sequestration Controllers

In this section, we evaluate how sensitive the steady-state error, stability margin, and robustness bound are to the process and controller degradation rates. Then, we provide guidelines for designing sequestration feedback controllers. Our goal is to design a robust sequestration feedback network with small steady-state error. As discussed in the previous Sections 2.1 and 2.2.2, the critical controller species degradation rate and the zero controller species degradation rate ensure zero steady-state error, but can be impractical to implement. To achieve a realistic and flexible design of sequestration feedback controllers, we relax the performance objective of zero steady-state error to the requirement of a small steady-state error. Using this performance objective instead, we demonstrate how to simultaneously tune the fragility and the relative steady-state error of a sequestration feedback network with two process species.

##### Example 1

(Find the Sequestration Controller for a Two Species Process Network). We consider the sequestration feedback network in Figure 4A and we design its sequestration controller. For simplicity, we assume that the process species production rates are equal such that *θ*_1_ = *θ*_2_ = *k*_1_ and that the process species degradation rates are also equal such that *γ_p_*_1_ = *γ_p_*_2_. Our requirements for optimal design include stability, robustness (a maximum disturbance amplification of at most two-fold), and a relative steady-state error of less than 30%.

First, we define the normalized process and controller rates as:

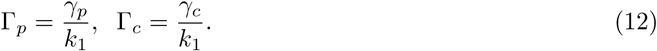

We note that the normalized process and controller rates, Γ*_p_* and Γ*_c_*, are unitless quantities, according to Table S1 and we use these normalized process and controller rates throughout this section. To assess stability, we define a rescaled empirical stability norm of a sequestration feedback network with two process species as:
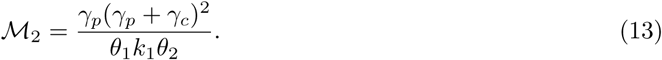

Then an empirical stability margin can be expressed as 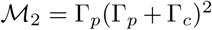. The stability criterion in equation (6) states that the sequestration feedback network is stable if and only if 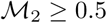.

Additionally, we express the relative steady-state error *ε*_2_ from the steady-state error *e*_2_ given in equation (11) in terms of the normalized process and controller species rates as follows:
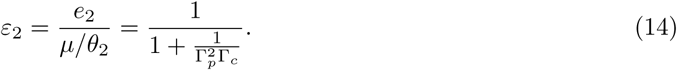

For simplicity, we also assume that we satisfy the first strong feedback assumption in Table 1, so that there is no critical controller species degradation rate that would result in a zero steady-state error. This assumption is equivalent to 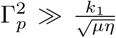. Moreover, to ensure that our sequestration controller design is not only stable, but also robust, we bound the infinity norm of its sensitivity function. In particular, a lower bound 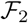 to the infinity norm of the sensitivity function can be analytically expressed using equation (7) as follows:
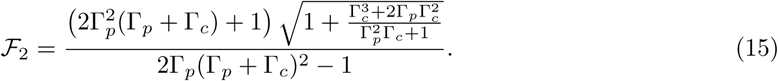

Now, we can simultaneously tune the lower bound 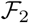 to the infinity norm of the sensitivity function and the relative steady-state error *ϵ*_2_ using Figure 6. A rule of thumb in engineering systems is that the magnitude of the infinity norm of the sensitivity function is between values 1.1 and 2 [Levine, 2010, Aström and Murray, 2008]. This implies that the maximum disturbance amplification through feedback is two-fold. Since we do not have an analytic expression for the sensitivity function itself, we instead bound 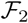 to be less than 2 in Figure 6A. This fragility value is indicated by the red dotted line in Figure 6A. Accordingly, the normalized process species degradation rate Γ*_P_* must be at least 0.7 and the normalized controller species degradation rate Γ*_C_* must be at least 0.8. We substitute values Γ*_P_* = 0.7 and Γ*_C_* = 0.8 in the relative steady-state error expression in equation (14) and compute a relative steady-state error of 28%. Thus, we fulfilled all the design specifications with a sequestration feedback network with values Γ*_P_* = 0.7 and Γ*_C_* = 0.8. Hence, the process species’ degradation rate must be 70% of the process species’ production rate and the controller species’ degradation rate must be 80% of the controller species’ production rate.

**Figure 6:**
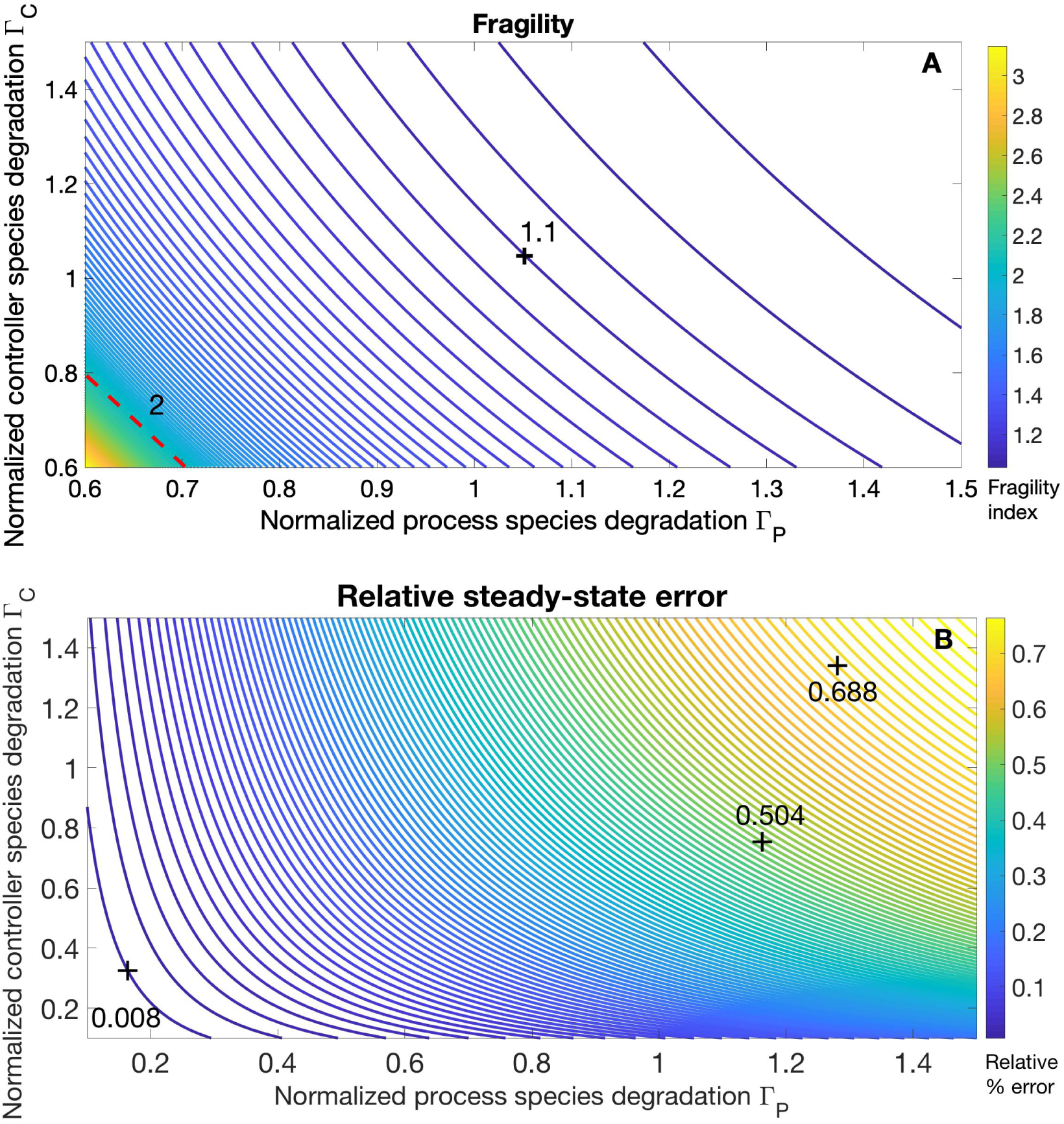
Guidelines for building robust controllers with small relative steady-state error. We plot both the fragility 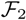 and the relative steady-state error *ε*_2_ as functions of the normalized process species’ degradation rate and the normalized controller species’ degradation rate. **A.** The red line indicates a value of 2, corresponding to a fragile system. Additional robustness can be achieved by increasing either the process or the controller species rates Γ*_p_* or Γ*_c_*. We note that all the feedback controllers in this panel are stable since 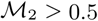, but some are fragile. **B.** The relative steady-state error approaches value zero for a smaller controller species degradation rate (integral control). As the process and the controller species’ degradation rates increase, the relative steady-state error approaches 68%. An extended plot of the fragility and relative steady-state error is available in the supplement (Figure S3). Using this plot, we note that both the fragility and the relative steady-state error functions are more sensitive to perturbations in the process species’ production and degradation rates than in the controller species’ production and degradation rates. This highlights the importance of measuring the process network’s rates for a robust, well-performing implementation of sequestration feedback.

To obtain these values, a biological implementation of the process and the controller species can be mRNA molecules or actively degraded proteins [Lewin, 2004]. The mRNA transcription rate more closely matches their degradation rate and thus can result in normalized process or controller degradation rates of 70% – 80% (Figure 5). Additionally, the rate of protein synthesis can also match the degradation rate, provided that the proteins are actively degraded by the proteosome. Sigma factors are a class of proteins that can also potentially result in normalized process or controller degradation rates of 70% – 80% (Figure 5).

##### Remark 1.

In Example 1, we assumed that the process species production rates and degradation rates were equal (*θ*_1_ = *θ*_2_ = *k*_1_ and *γ_p1_* = *γ_p2_*). These assumptions served to simplify the expressions in equations (7) and (11). More generally, the sequestration controller design for a network with two process species is analogous to the design in Example 1, provided that the strong feedback assumptions hold. The strong feedback assumptions are explicitly given in Theorem S1. For controller design, equations (8) and (11) can be used to modulate the stability margin and steady-state error of the sequestration network.

#### 2.3.3 Large Stability Margin and Small Steady-State Error Are Competing Objectives

Achieving a large stability margin and a small steady-state error are competing objectives for sequestration feedback networks, as noted in [Baetica, 2018, Olsman et al., 2018] and illustrated in Figure 6. To achieve a small relative steady-state error, both the normalized process and controller species degradation rates (Γ*_p_* and Γ*_c_*) should be as small as possible. The relative steady-state error is a monotonically increasing function of both of these rates. This property can be observed both from equation (14) and from Figure 6B. However, to achieve a large stability margin (or robustness), both the normalized process and controller species degradation rates (Γ*_p_* and Γ*_c_*) should be as large as possible. This property can be observed both from equations (13) and (15), as well as from Figure 6A. Therefore, attaining a large stability margin and a small steady-state error are competing design objectives. It is important to note that these two competing objectives can be particularly challenging to achieve for process networks with small and large ratios between their degradation and their production rates.

The motivation for studying cases of extreme competition between these two objectives is that in practice, process networks are often already specified or challenging to modify. Consequently, the process species’ production and degradation rates are often fixed. When the process species’ degradation rate is much smaller than the process species’ production rate, then the parameter Γ*_p_* can be small. Thus, a large controller species degradation rate is required to ensure that stability is achieved and an even larger controller species degradation rate is required to guarantee robustness. For a small value of Γ*_p_*, it must be that
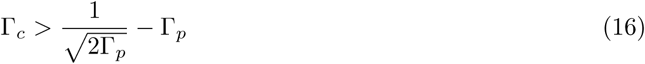

simply to achieve stability and an even larger controller degradation rate for robustness. For example, for a process network with value of Γ*_p_* = 0.1 and a desired a fragility value of 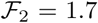, the normalized controller species rate Γ*_c_* must be at least 12.5. In this case, the relative steady-state error is approximately 11%. To achieve a robust controller design for a slow process network, we recommend implementing the controller species using RNA or actively degraded protein parts.

When the process species degradation rate is much larger than its production rate, we obtain large values of Γ*_p_*. This corresponds to actively degraded proteins and sigma factors or to proteins with high production rates. In this case, the fragility function approaches value one, whereas the steady-state error needs to be bounded such that
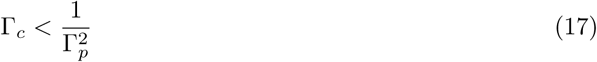

to obtain a 50% relative steady-state error. To further reduce the relative steady-state error, the controller species degradation rate must be even smaller. Here, it is more advantageous to have a very small controller species degradation rate. We recommend an implementation that uses slowly degrading proteins for the sequestration controller so as to reduce the magnitude of the relative steady-state error.

Finally, we note that the process species’ production and degradation rates have a bigger impact on the steady-state error and the fragility functions than the controller species’ production and degradation rates (Figure S3). The impact of the process species’ production and degradation rates highlights the importance of measuring these rates before building a sequestration controller for the network. Depending on the process network’ properties, the competition between achieving stability, robustness, and small relative steady-state error can be more or less severe. For particularly severe cases, we have provided guidelines for the design of the controller species in equations (16) and (17), along with suggestions for the corresponding biological parts.

## 3 Discussion

The development of a first generation of biological controllers for bacteria and yeast marks the beginning of an era when synthetic biological systems can function robustly and perform well. In this work, we have considered synthetic biological controllers implemented by a class of sequestration feedback networks. Using control theoretical methods, we have proposed designs that ensure stability, robustness, and good performance for these sequestration feedback network. We have offered tuning options for the strength of the controller species’ sequestration reaction, as well as for the production and the degradation rates of both the process and the controller species. When possible, we have suggested biological parts for the practical implementations of these designs.

The application of control theory to synthetic biological controllers aims to ensure that they function robustly, in different host organisms, despite perturbations in their environments. Nevertheless, almost all sequestration feedback controllers in the current synthetic biology literature have been concurrently built with the process networks they control [McCardell et al., 2017, Hsiao et al., 2014, Folliard, 2017, Lillacci et al., 2017, Chevalier et al., 2018]. This approach will likely limit the versatility of these biological controllers as they will be optimal with respect to a single process network, a single host organism, and its environment. Therefore, we believe that it is important to continue assessing the realistic constraints imposed by building sequestration feedback controllers for fixed process networks, as we have done in Section 2.3.3. By incorporating realistic constraints in our design of sequestration feedback controllers, we guarantee that they function robustly, across a variety of environmental conditions and biological host organisms.

Moreover, the engineering design tools introduced in this paper are applicable to other mechanisms for synthetic biological control, in addition to sequestration feedback. Several mechanisms for biological control that are currently being explored include paradoxical extracellular signaling [Hart et al., 2014] and post-translation mechanisms such as multi-protease regulation. Other biochemical reaction network designs that implement robust perfect adaptation through integral control have been proposed in [Xiao and Doyle, 2018]. Using similar control theoretical tools, it is possible to develop models for these biological controllers and to assess their properties of stability, robustness, and performance. Depending on the applications of interest to synthetic biology, we will benefit from multiple mechanisms for feedback control of synthetic systems and from multiple feedback controller designs.

## Author Contributions

Conceptualization and Methodology, AAB, YPL, NO, and RMM; Formal Analysis, AAB, YPL, and NO; Software, AAB and YPL; Writing, AAB; Supervision and Funding, RMM.

## Acknowledgements

The authors would like to thank Michael W. Chevalier and Hana El-Samad for providing feedback on the manuscript. The project was sponsored by the Defense Advanced Research Projects Agency (Agreement HR0011-17-2-0008). The content of the information does not necessarily reflect the position or the policy of the Government, and no official endorsement should be inferred.

## Declaration of Interests

The authors declare no competing interests.

## S4 Supplemental Information

### S4.1 The Properties Of The Sequestration Controller

In this section, we demonstrate that the sequestration controller implements integral feedback (perfect adaptation) when the two controller species are not subjected to degradation, provided that the closed loop sequestration network is stable. The property of perfect adaptation results in a zero steady-state error for the sequestration feedback network in Figure 2. We also demonstrate that when the controller species are subjected to degradation and dilution, the sequestration controller is a lag compensator. Moreover, we analyze the properties of the steady-state error of sequestration feedback with nonzero controller species degradation in Section S4.3. We give criteria for the stability of sequestration feedback networks in [Olsman et al., 2018, Baetica, 2018].

When the controller species are not degraded or diluted, the model of their dynamics is as follows:
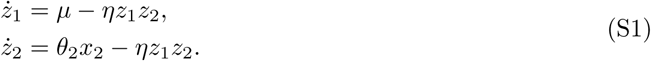

We can simply subtract the two differential equations in equation (S1) to obtain that
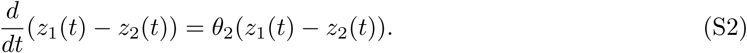

Assuming that the reference signal has value 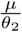, then we can define the error signal as 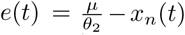. Thus, the controller species implement integral control since the control action *z*_1_(*t*) − *z*_2_(*t*) integrates the error signal *e*(*t*) as follows:
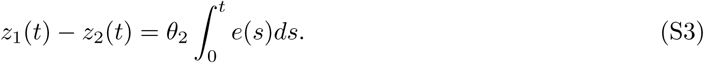

If the sequestration feedback network is stable, then the model of its dynamics has a steady-state. At steady-state, the integral controller ensures that the property of perfect adaptation holds since
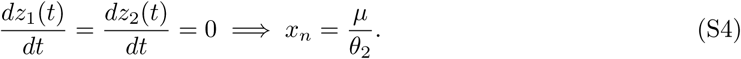

Perfect adaptation is a desirable property of the sequestration feedback system because it allows for a variety of process network dynamics. We now investigate whether perfect adaptation is retained when we include the controller species degradation and dilution in the model. By incorporating the degradation of the controller species in the model description in equation (S1), the controller dynamics each gain an additional term:
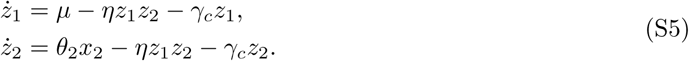

As before, we subtract the two equations that describe the controller dynamics in equation (S5). From this subtraction, we infer that the resulting sequestration controller is a lag compensator. The compensator integrates the error signal weighed by an exponential of the controller degradation rate as follows:
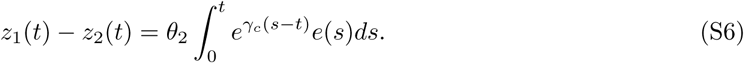

Both the integral controller and the lag compensator act as filters for the past error signal. Intuitively, the integral controller integrates the error signal with uniform weight, whereas the lag compensator integrates the error signal with bias towards more recent error measurements. This occurs because the exponential of the degradation rate biases the error measurement towards recent past over the distant past since 0 ≤ *s* ≤ *t* in equation (S6).

More importantly, the lag compensator has different properties than integral control in terms of the magnitude of the steady-state error. Since integral control has the property of perfect adaptation, provided that the closed loop system in stable, then the closed loop system has zero steady-state error. The lag compensator can also exhibit zero steady-state error for a specific controller degradation rate, as demonstrated in Section S4.3, but otherwise results in steady-state error.

### S4.2 Alternative Modeling Assumptions

In this section, we consider different modeling assumptions than the setup introduced in Figure 3. Firstly, we assume that the process species in the sequestration feedback network degrade at different rates. If the process species are degraded at different rates, the characteristic polynomial in [Olsman et al., 2018] does not necessarily factor the term (*s* + 1)*^n^* for *n* > 2 process species. Therefore, finding an analytic approximation to the roots of the characteristic polynomial can be challenging and finding an analytic criterion for the stability of sequestration feedback networks may not be possible.

However, when there are only two process species as in Figure 4A, we can find an analytical criterion for stability under strong sequestration feedback assumptions. As indicated by the simulation in Figure 4C, the smaller of the two process species degradation rates results in instability. Although the larger process species degradation rate dampens the oscillations (compare panels B and D), it is not able to fully compensate for the small degradation rate of the other process species. Therefore, the smallest process species degradation rate must be carefully tuned to ensure stability of the sequestration feedback network. We model the effect of the two different process species degradation rates *γ*_*p*1_ and *γ*_*p*2_ as follows:
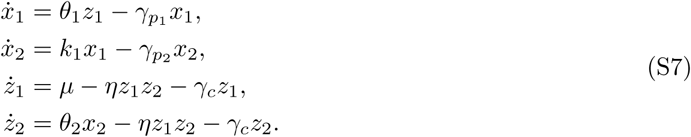

#### Theorem S1

(Stability Criterion for a Sequestration Feedback Network with Two Process Species). *We consider the sequestration feedback network in Figure 4A with the dynamics given in equation (S7). If this sequestration network fulfills the strong feedback assumptions*
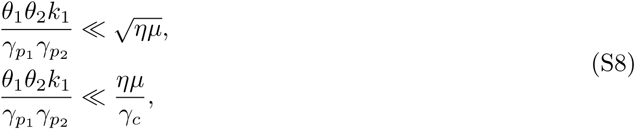

*then the closed loop sequestration feedback network is stable if and only if*
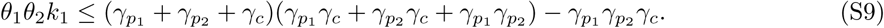

*Proof.* When the sequestration feedback network only has two process species, we can derive an analytical criterion for its stability. In this case, the characteristic polynomial associated to the linearized sequestration feedback network is:
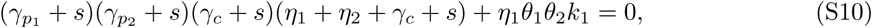

where 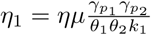 and 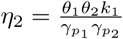 for complex roots 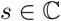. Under the strong sequestration feedback assumptions, the characteristic polynomial simplifies to:
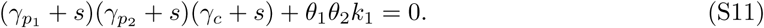

For a purely complex root *s* = *iω*, we set the both the real and the imaginary parts of the polynomial in equation (S11) to zero. Hence, it must be that
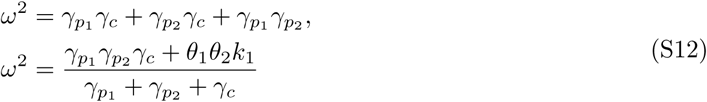
simultaneously. Therefore, according to Descartes’ rule of sign, the sequestration feedback network in Figure 4A is stable under the strong feedback assumptions if and only if
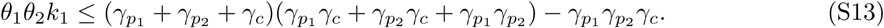

An alternative set of modeling assumptions is to consider the reactions between the process and the controller networks to be enzymatic instead of catalytic. We note that the catalytic reactions between the process and the controller networks in Figure 3 can be described as:
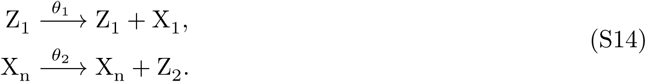

Alternatively, we can assume that the transformation of controller species *Z*_1_ into the process input species *X*_1_ is catalyzed by the enzyme *E*_1_ and that the transformation of the process output species *X_n_* into the controller species *Z*_2_ is catalyzed by the enzyme *E*_2_. Then the dynamics of these catalytic reactions can be modeled using the Michaelis-Menten kinetics as follows:
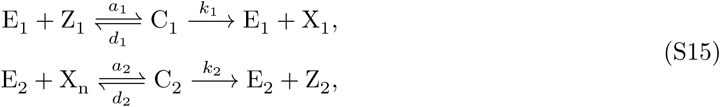

where the rates satisfy the inequalities *k*_1_ ≫ *a*_1_, *d*_1_ and *k*_2_ ≫ *a*_2_, *d*_2_. We make the simplifying assumptions that the concentration of enzymes *E*_1_ and *E*_2_ are much smaller than the concentrations of species *Z*_1_ and *X*_1_ [Mathews et al., 2000, Del Vecchio and Murray, 2015]. We let the Michaelis constants be
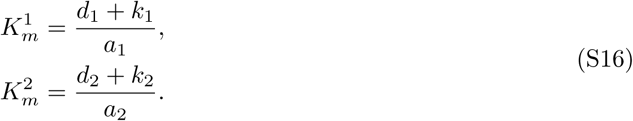

The Michaelis constants 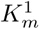 and 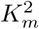 that correspond to biological enzymes range between 10^−6^ M and 10^−2^ M [Mathews et al., 2000], which renders them to be much larger than the concentrations of species *Z*_1_ and *X_n_*. Therefore, the propensity functions associated with these kinetics are linear.

We let 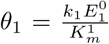 and 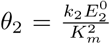, where 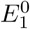 and 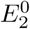 are the initial concentrations of enzymes *E*_1_ and *E*_2_, respectively. Using this notation, the sequestration feedback model remains unchanged from Figure 3 due to the linearity of the first-order kinetics. However, if the simplifying assumptions that the concentration of enzymes *E*_1_ and *E*_2_ are much smaller than the concentrations of species *Z*_1_ and *X*_1_ do not hold, then the sequestration feedback controller may not behave as a lag compensator, as suggested in [Xiao and Doyle, 2018].

### S4.3 The Critical Controller Species Degradation Rate

In the previous sections, we derived analytic criteria for the stability of the sequestration feedback networks. Subsequently, we analyze their performance properties and in particular, we focus on the magnitude of the steady-state error. We have previously stated that the closed loop system can have zero steady-state error for no controller species degradation, as well as for a value of the controller degradation rate, which we refer to as “critical” (Figure S1B). Assuming that the sequestration feedback network is stable, a zero controller species degradation guarantees perfect adaptation of the closed loop system, but can be challenging to implement. In this section, we analytically derive a value of the critical controller species degradation rate such that the steady-state error of a more general stable sequestration feedback network with *n* process species equals zero. We also demonstrate that the degradation rate values of zero and of the critical degradation rate are the only ones for which the network tracks the reference with zero steady-state error. Depending on the parameters of the sequestration feedback system, the critical value of the degradation rate may or may not be achievable.

We consider a deterministic sequestration feedback network with *n* process species as in Figure 2. However, we restrict the dynamics of the *n* process species to be linear (the process species can be degraded or can serve as catalysts for the creation of other process species). We restrict our analysis to linear process species dynamics because otherwise the analytical derivation of the linearization around steady-state is challenging. Then the model of the sequestration feedback network is given by the following system of equations:
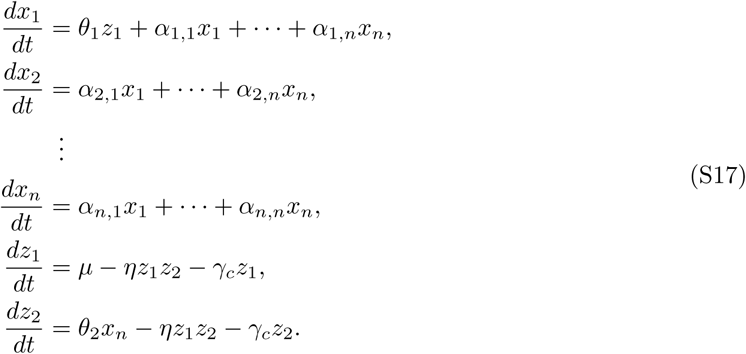

We define the following notation:
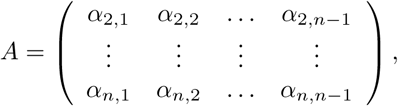

*α*_1_ = (*α*_1,1_, …, *α*_1,*n*−1_), *α_n_* = (*α*_2,*n*_, …, *α_n,n_*)*^T^*, and 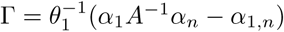.

#### Theorem S2

(The Critical Controller Degradation Rate). *The critical controller degradation rate of the sequestration feedback network with n process species modelled in equation (S17) is given by:*
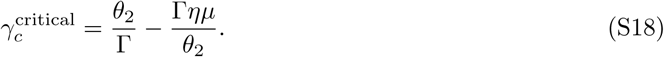

*The critical controller species degradation rate exists if and only if the closed loop system is stable and* 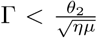, *where* 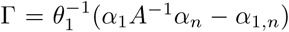. *Otherwise, the steady-state error is nonzero for all nonzero controller species degradation rates.*

**Figure S1:**
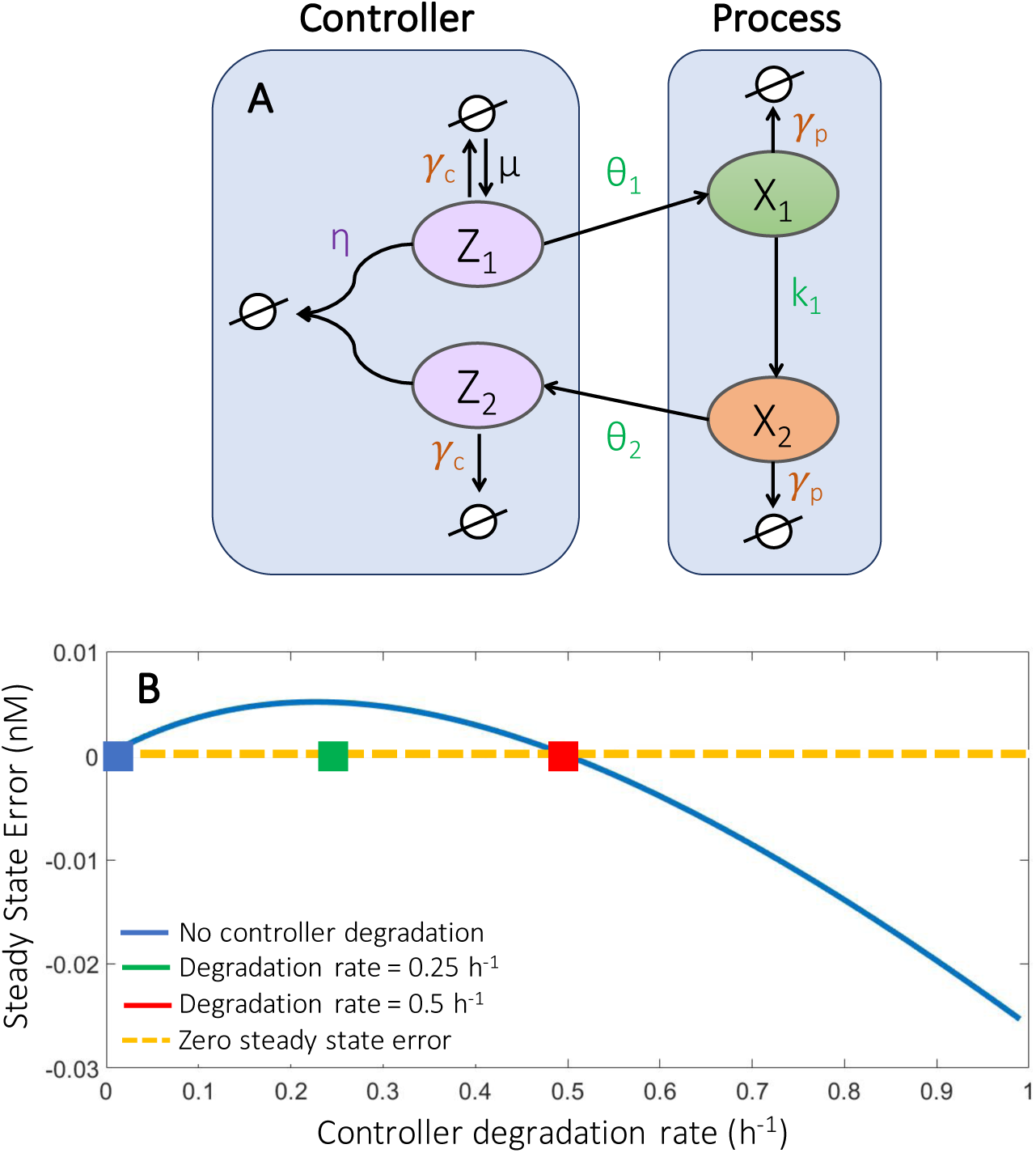
The steady-state error as a function of the controller degradation rate. **A.** We introduce a sequestration feedback network with two process species. For the numerical simulations in this section and in Section S4.4, we consider the sequestration feedback network with only two process species *X*_1_ and *X*_2_. We assume that process species *X*_1_ and *X*_2_ are degraded at the same rate *γ_p_* and that the controller species *Z*_1_ and *Z*_2_ are degraded at the same rate *γ_c_*. We restate the model in equation (S33). **B.** We plot the steady-state error (blue line) as a function of the controller degradation rate, while keeping the other parameters of the sequestration network in panel A fixed. We obtain perfect adaptation for no controller degradation (blue square) and zero steady-state error at the critical controller degradation rate (red square). The steady-state error is nonzero at other controller species degradation rates (green square). Zero and the critical controller species degradation rate are the only two degradation rate values for which the steady-state error equals zero (orange dashed line). This property holds for a more general class of sequestration feedback networks, as discussed in Theorem S2. We derive an analytic expression for the steady-state error as a function of the controller species degradation rate in equation (S23).

*Proof.* In a sequestration feedback system with a single process species, the critical degradation rate
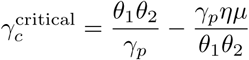
results in zero steady-state error. This degradation rate value can only be achieved when 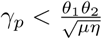.

We now consider a sequestration feedback network with *n* > 1 process species. Assuming that the closed loop system is stable, at equilibrium, equation (S17) reduces to
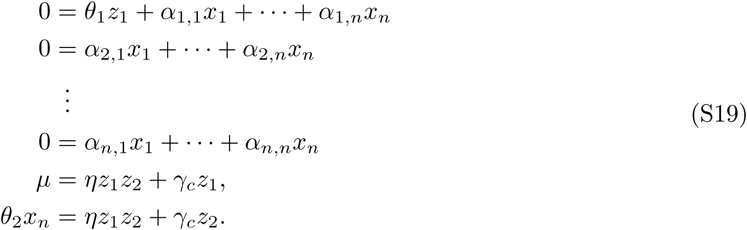

Using the notation introduced in this section, the system of equations in (S19) is equivalent to
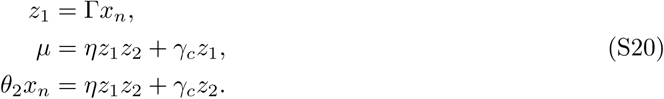

First, it must be the case that constant Γ > 0 and matrix *A* is invertible. Otherwise, the system cannot have a positive steady-state. This is equivalent to *α*_1_*A*^−1^*α_n_* > *α*_1,*n*_. The input species should not be depleted to create the output species. The system in equation (S20) simplifies to a single equation
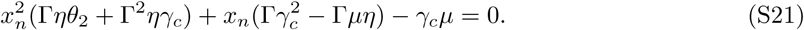

Equation (S21) always has a positive solution
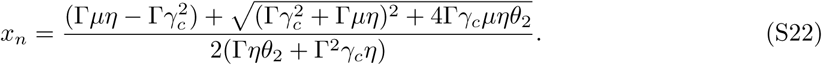

Thus, the steady-state error signal is
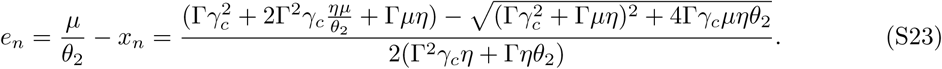

If we want the output of the dynamical system to follow the reference signal 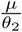, then it must be that
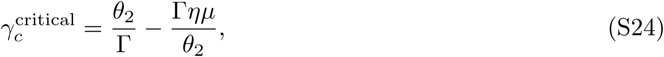
which can only be achieved if and only if 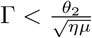.

#### Theorem S3

(The Critical Controller Degradation Rate Exists only under Particular Conditions). *The critical controller degradation rate for a simplified process network (i.e. each process species X_i_ is created by the previous process species X*_*i* − 1_ *and creates the next process species X_i_*_+1_, 2 ≤ *i* ≤ *n* − 1, *as in Figure 3) is:*
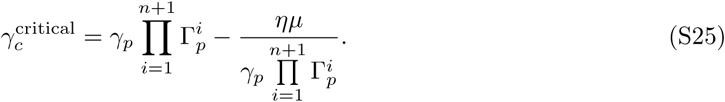

*For this sequestration feedback network, the critical controller species degradation rate exists if and only if the process species degradation rate satisfies the inequality*
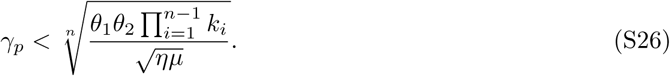

*Proof.* The dynamics of this sequestration feedback network can be modelled as follows:
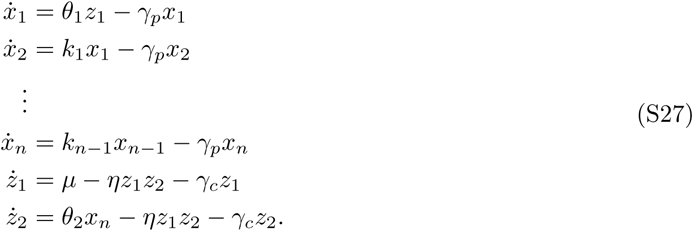

Therefore, the simplified matrices for the dynamics of the network are:
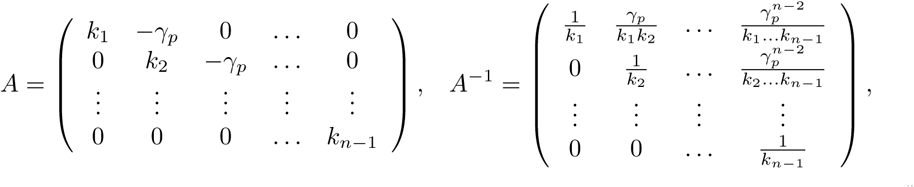

the vectors *α*_1_ = (−*γ_p_*, 0, …,0), *α_n_* = (0, …,0, −*γ_p_*)*^T^*, *α*_1, *n*_ = 0, and the expression 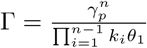.

According to Theorem S2, the critical controller degradation rate is
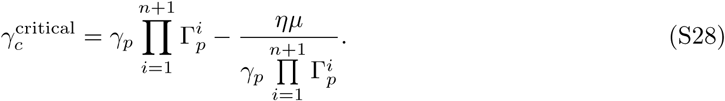

The critical controller degradation rate exists only when
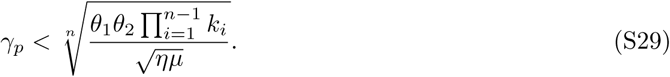

This condition is equivalent to equation (10) in the main text of the paper. The analytic expression of the error function for this particular class of sequestration networks is
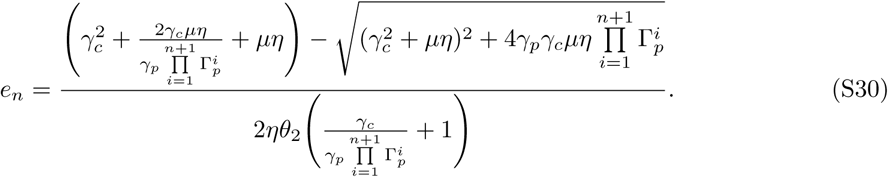

#### Remark S2.

We note that the strong feedback assumptions imply that the critical controller degradation rate does not exist. Under the strong feedback assumptions in Table 1, the analytic expression of the steady-state error simplifies to
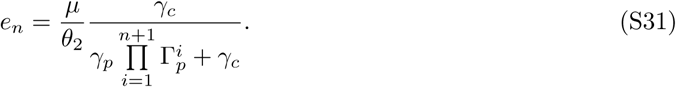

Thus, the steady-state error is a monotonically increasing function of the controller species degradation rate. We refer to the ratio between the steady-state error and the reference value as the relative error. Using this simplified expression, the relative steady-state error function can be bounded. For example, if we are interested in obtaining a relative steady-state error of less than 10%, then we can design a controller degradation rate such that
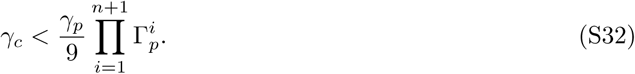

### S4.4 Additional Simulation Details

#### S4.4.1 Scaling the sequestration feedback network model

To produce the simulations in Figures S1B and in Sections S4.2 and S4.4, we first rescale the model of the sequestration feedback network. For simplicity, we only describe the rescaling step for a process network with two species (Figure S1A). However, this rescaling step is applicable for a general linear process network. The unscaled model of the sequestration feedback network is:
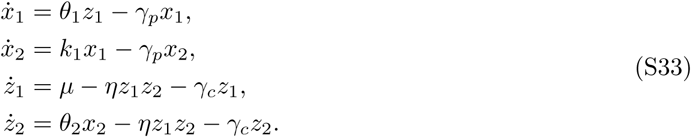

**Table S1:** The units of the biochemical species and rates in the unscaled and scaled sequestration feedback network. Species *X*_1_, *X*_2_, *Z*_1_, and *Z*_2_ have units of molar (M) and time t has units of seconds (s). Thus, the degradation and the production rates *θ*_1_, *θ*_2_, *γ_p_*, *γ_c_*, and *k*_1_ have units of s^−1^ since they correspond to first order reactions. Rate *µ* has units of M s^−1^ since it corresponds to a zero order reaction. Rate *η* has units of M^−1^ s^−1^ since it corresponds to a second order reaction. Following rescaling, biochemical species 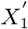, 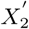, 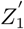, and 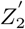 have units of nanomolar (nM) and time t′ has units of hours (h). Thus, the degradation and the production rates 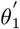, 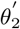, 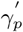, 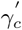, and 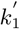 have units of h^−1^ since they correspond to first order reactions. Rate *µ*′ has units of nM h^−1^ since it corresponds to a zero order reaction. Rate *η*′ has units of nM^−1^ h^−1^ since it corresponds to a second order reaction.

|  |  |  |  |  |  |  |  |  |  |
| --- | --- | --- | --- | --- | --- | --- | --- | --- | --- |
| unscaled | $x_1, x_2, z_1, z_2$ | t | $\theta_1$ | $\theta_2$ | $\gamma_p$ | $\mu$ | $\eta$ | $\gamma_c$ | $k_1$ |
| | M | s | $\text{s}^{-1}$ | $\text{s}^{-1}$ | $\text{s}^{-1}$ | $\text{M s}^{-1}$ | $\text{M}^{-1} \text{s}^{-1}$ | $\text{s}^{-1}$ | $\text{s}^{-1}$ |
| scaled | $x'_1, x'_2, z'_1, z'_2$ | t' | $\theta'_1$ | $\theta'_2$ | $\gamma'_p$ | $\mu'$ | $\eta'$ | $\gamma'_c$ | $k'_1$ |
| | nM | h | $\text{h}^{-1}$ | $\text{h}^{-1}$ | $\text{h}^{-1}$ | $\text{nM h}^{-1}$ | $\text{nM}^{-1} \text{h}^{-1}$ | $\text{h}^{-1}$ | $\text{h}^{-1}$ |

The units of the unscaled biochemical species and rates are given in Table S1. To rescale, we let 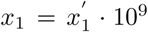, 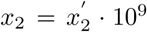, 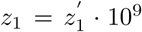, 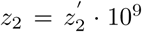, 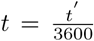, 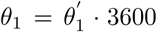, 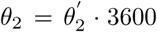, 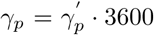, *γ_c_ = γ_c_* · 3600, 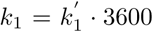, *η* = *η*′ · 10^−9^ · 3600, and *µ* = *µ*′ · 10^9^ · 3600. Following rescaling, we obtain new units of the biochemical species and rates (Table S1).

Following rescaling, the model of the sequestration feedback network can be described as:
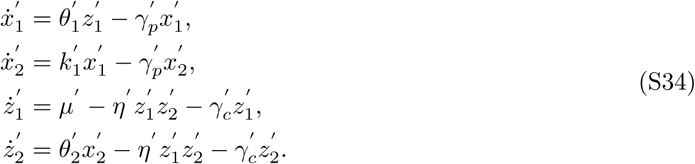

**Figure S2:**
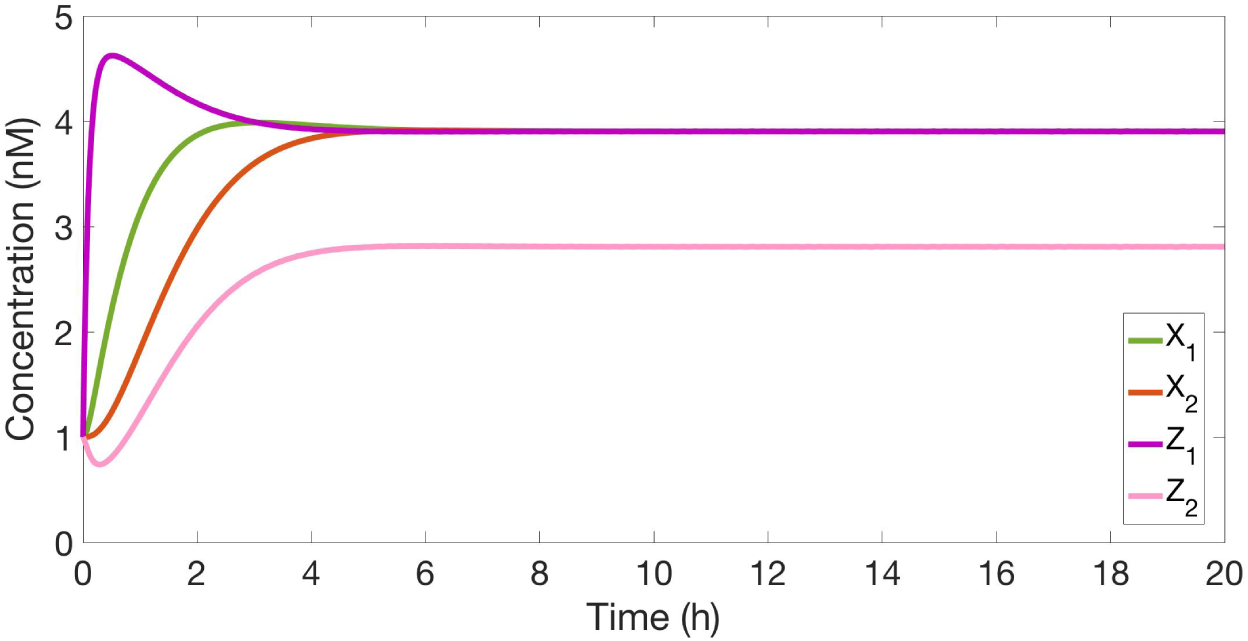
The sequestration feedback network with the controller species implemented with transcriptional parts and the process species implemented with protein parts. The sequestration feedback network simulated using the parameters: 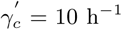, *η*′ = 1 nM^−1^, 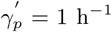, *µ*′ = 50 nM h^−1^, 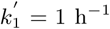, 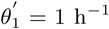, 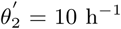. The values of parameters *γ_p_*, *γ_c_*, and *η* are found in Figure 5 for both the transcriptional and the protein parts. The sequestration feedback network is stable and has oscillations that settle within 5 hours. The reference concentration is 5 nM and the steady-state error is 1.1 nM.

Following the rescaling step, we use representative values of the reaction rates in equation (S34) to perform simulations and gain intuition about the biological implementation of sequestration feedback networks. We compute representative values of the controller species’ and the process species’ degradation rates in Figure 5 in units of h^−1^ by taking the inverses of the half-life values and multiplying them by the constant log(2). The median mRNA half-life is measured as 0.5-6 min in [Bernstein et al., 2004]. In the type I toxin-antitoxin pair in *E. coli*, the half-life of toxin Hok is 20 minutes [Steif and Meyer, 2012]. The half-life of the sigma factor protein RpoS is 30 min when the *E. coli* cells are in stationary phase at 37°C or under stress conditions. When sigma factor proteins RpoS are actively degraded by protease ClpXP during the exponential phase, their half-life is only 2 minute [Zhou and Gottesman, 1998]. The antitoxin CcdA is degraded in wild-type cells with a half-life of 30 min in the absence of toxin CcdB and a half-life of 60 min when bound in a complex with toxin CcdB [De Jonge et al., 2009]. Using these half-life measurements, we compute the degradation rate values in Figure 5.

Lastly, we give representative values of the sequestration binding on-rates in Figure 5. The rate *η*′ in the rescaled sequestration feedback model (equation (S34)) represents the on-rate of the sequestration reaction between the two controller species. The off-rate of the sequestration reaction is much smaller than the on-rate. To match the units in Table S1, we give the on-rates of the sequestration reactions in units of nM^−1^ h^−1^. From [Walton et al., 2002], we compute the representative on-rate values of mRNA binding to antisense RNA to be between 0.005 nM^−1^ h^−1^ and 1.62 nM^−1^ h^−1^. We compute representative on-rate values of sigma factors binding anti-sigma factors between 18 nM^−1^ h^−1^ and 72 nM^−1^ h^−1^, depending on the temperature and the presence of zinc from [Sharma and Chatterji, 2008, Rajasekar et al., 2016]. To the best of our knowledge, the on-rate of the toxin CcdB binding the antitoxin CcdA has not been accurately determined [Kampranis et al., 1999].

#### S4.4.2 Details of numerical experiments

For the two-species process network in Figure S1A, both quantities 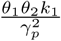 and *ηµ* have units of h^−1^. For a transcriptional implementation of the controller species and a protein implementation of the process species, this network satisfies the assumption of strong sequestration feedback for the parameter values in Figure S2.

For the design example in Section 2.3.2, we plot the fragility 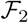 and relative steady-state error *ε*_2_ as functions of the normalized process and controller species degradation rates (Γ*_p_* and Γ*_c_*) for a more extensive range of parameters than in Figure 6. The more extensive range of parameter values enables us to observe that the contour plots are asymmetric, demonstrating more sensitivity towards the process species rate Γ*_p_*. Hence, the process network’s parameters have more impact on fragility and steady-state error than the controller network’s parameters. Thus, it is paramount to obtain measurements of the process network’s parameters for a successful design of sequestration control.

**Figure S3:**
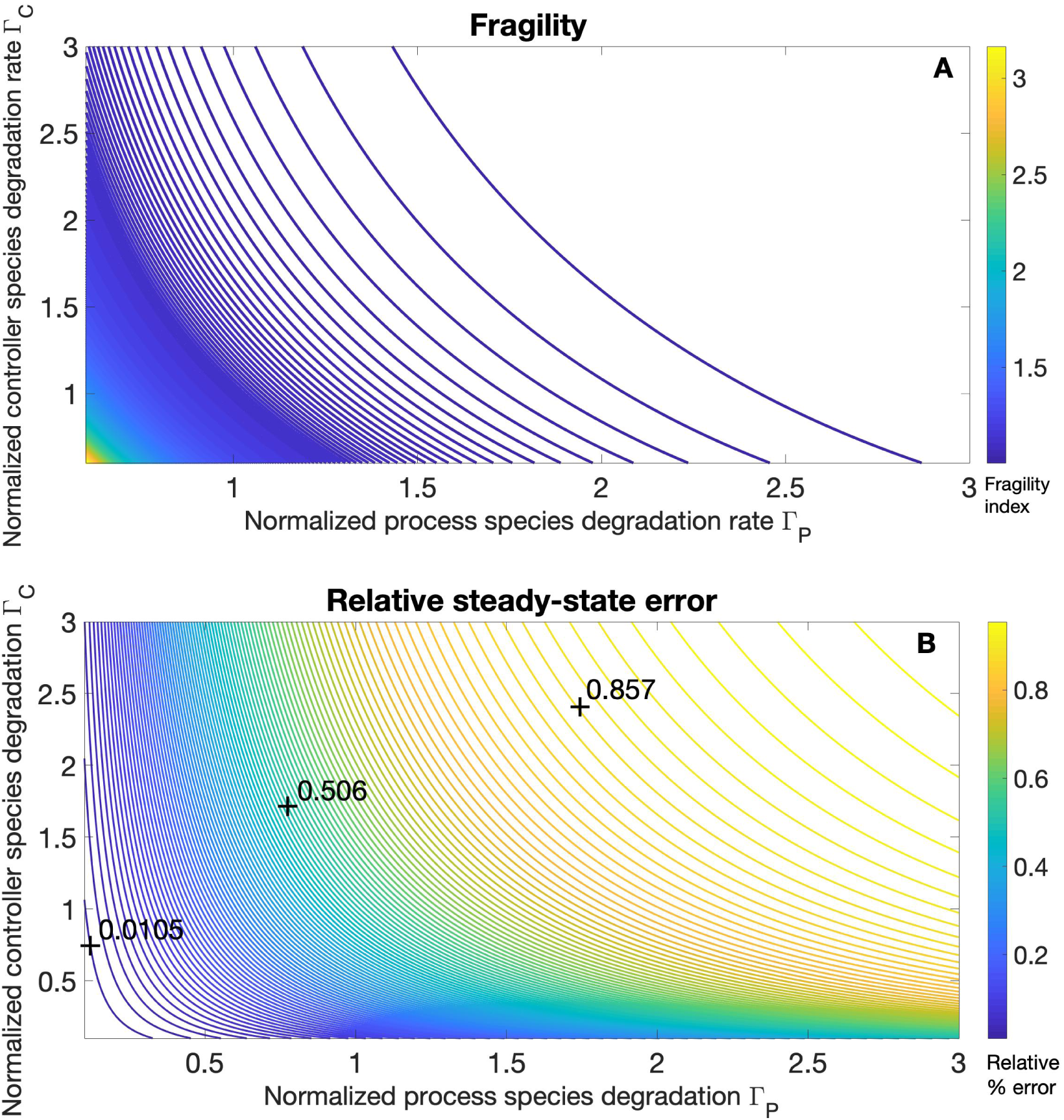
Both fragility and relative steady-state error are more sensitive to the parameters of the process network than to the parameters of the controller network. We plot the fragility 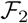 and the relative steady-state error *ε*_2_ as functions of the normalized process and controller species degradation rates (Γ*_p_* and Γ*_c_*) for a more extensive range of parameters than in Figure 6. As the process and the controller species degradation rates increase, the fragility index approaches value three and the relative steady-state error approaches value one. Since the contour functions are asymmetric, we infer that the fragility and relative steady-state functions are more sensitive to perturbations in the process species’ production and degradation rates than in the controller species’ production and degradation rates. This highlights the importance of measuring the process network’s production and degradation rates when designing its sequestration controller.

## References

[Ang et al., 2010] Ang, J., Bagh, S., Ingalls, B. P., and McMillen, D. R. (2010). Considerations for using integral feedback control to construct a perfectly adapting synthetic gene network. Journal of theoretical biology, 266(4):723–738.

[Annunziata et al., 2017] Annunziata, F., Matyjaszkiewicz, A., Fiore, G., Grierson, C. S., Marucci, L., di Bernardo, M., and Savery, N. J. (2017). An orthogonal multi-input integration system to control gene expression in escherichia coli. ACS Synthetic Biology, 6(10):1816–1824.

[Arkin, 2008] Arkin, A. (2008). Setting the standard in synthetic biology. Nature Biotechnology, 26(7):771.

[Arnold, 1998] Arnold, F. H. (1998). Design by directed evolution. Accounts of Chemical Research, 31(3):125–131.

[Aström and Murray, 2008] Aström, K. J. and Murray, R. M. (2008). Feedback Systems: An Introduction For Scientists And Engineers. Princeton University Press.

[Baetica, 2018] Baetica, A.-A. (2018). Design, Analysis, And Computational Methods For Engineering Synthetic Biological Networks. PhD thesis, California Institute of Technology.

[Bernstein et al., 2002] Bernstein, J. A., Khodursky, A. B., Lin, P.-H., Lin-Chao, S., and Cohen, S. N. (2002). Global analysis of mrna decay and abundance in escherichia coli at single-gene resolution using two-color fluorescent dna microarrays. Proceedings of the National Academy of Sciences, 99(15):9697–9702.

[Bernstein et al., 2004] Bernstein, J. A., Lin, P.-H., Cohen, S. N., and Lin-Chao, S. (2004). Global analysis of escherichia coli rna degradosome function using dna microarrays. Proceedings of the National Academy of Sciences of the United States of America, 101(9):2758–2763.

[Briat et al., 2016] Briat, C., Gupta, A., and Khammash, M. (2016). Antithetic integral feedback ensures robust perfect adaptation in noisy biomolecular networks. Cell Systems, 2(1):15–26.

[Canton et al., 2008] Canton, B., Labno, A., and Endy, D. (2008). Refinement and standardization of synthetic biological parts and devices. Nature Biotechnology, 26(7):787.

[Chandra et al., 2011] Chandra, F. A., Buzi, G., and Doyle, J. C. (2011). Glycolytic oscillations and limits on robust efficiency. science, 333(6039):187–192.

[Cherny and Gazit, 2008] Cherny, I. and Gazit, E. (2008). Amyloids: Not only pathological agents but also ordered nanomaterials. Angewandte Chemie International Edition, 47(22):4062–4069.

[Chevalier et al., 2018] Chevalier, M., Gomez-Schiavon, M., Ng, A., and El-Samad, H. (2018). Design and analysis of a proportional-integral-derivative controller with biological molecules. bioRxiv.

[Clomburg and Gonzalez, 2010] Clomburg, J. M. and Gonzalez, R. (2010). Biofuel production in escherichia coli: The role of metabolic engineering and synthetic biology. Applied Microbiology and Biotechnology, 86(2):419–434.

[Dahiyat and Mayo, 1996] Dahiyat, B. I. and Mayo, S. L. (1996). Protein design automation. Protein Science, 5(5):895–903.

[Daniel et al., 2013] Daniel, R., Rubens, J. R., Sarpeshkar, R., and Lu, T. K. (2013). Synthetic analog computation in living cells. Nature, 497(7451):619.

[De Jonge et al., 2009] De Jonge, N., Garcia-Pino, A., Buts, L., Haesaerts, S., Charlier, D., Zangger, K., Wyns, L., De Greve, H., and Loris, R. (2009). Rejuvenation of ccdb-poisoned gyrase by an intrinsically disordered protein domain. Molecular cell, 35(2):154–163.

[Del Vecchio, 2015] Del Vecchio, D. (2015). Modularity, context-dependence, and insulation in engineered biological circuits. Trends in biotechnology, 33(2):111–119.

[Del Vecchio and Murray, 2015] Del Vecchio, D. and Murray, R. M. (2015). Biomolecular feedback systems. Princeton University Press.

[DeLoache et al., 2015] DeLoache, W. C., Russ, Z. N., Narcross, L., Gonzales, A. M., Martin, V. J., and Dueber, J. E. (2015). An enzyme-coupled biosensor enables (s)-reticuline production in yeast from glucose. Nature chemical biology, 11(7):465.

[Doyle et al., 2013] Doyle, J. C., Francis, B. A., and Tannenbaum, A. R. (2013). Feedback control theory. Courier Corporation.

[Dunlop et al., 2010] Dunlop, M. J., Keasling, J. D., and Mukhopadhyay, A. (2010). A model for improving microbial biofuel production using a synthetic feedback loop. Systems and Synthetic Biology, 4(2):95–104.

[El-Samad et al., 2002] El-Samad, H., Goff, J., and Khammash, M. (2002). Calcium homeostasis and parturient hypocalcemia: An integral feedback perspective. Journal of Theoretical Biology, 214(1):17–29.

[Elowitz and Leibler, 2000] Elowitz, M. B. and Leibler, S. (2000). A synthetic oscillatory network of transcriptional regulators. Nature, 403(6767):335.

[Ferrell Jr, 2016] Ferrell Jr, J. E. (2016). Perfect and near-perfect adaptation in cell signaling. Cell Systems, 2(2):62–67.

[Folliard, 2017] Folliard, T. (2017). A synthetic viral feedback loop results in robust expression. ACS Synthetic Biology.

[Friedland, 2012] Friedland, B. (2012). Control system design: an introduction to state-space methods. Courier Corporation.

[Gardner et al., 2000] Gardner, T. S., Cantor, C. R., and Collins, J. J. (2000). Construction of a genetic toggle switch in escherichia coli. Nature, 403(6767):339–342.

[Georgianna and Mayfield, 2012] Georgianna, D. R. and Mayfield, S. P. (2012). Exploiting diversity and synthetic biology for the production of algal biofuels. Nature, 488(7411):329.

[Gerdes, 1988] Gerdes, K. (1988). The parb (hok/sok) locus of plasmid r1: a general purpose plasmid stabilization system. Nature Biotechnology, 6(12):1402–1405.

[Goentoro et al., 2009] Goentoro, L., Shoval, O., Kirschner, M. W., and Alon, U. (2009). The incoherent feedforward loop can provide fold-change detection in gene regulation. Molecular Cell, 36(5):894–899.

[Gomez et al., 2016] Gomez, M. M., Murray, R. M., and Bennett, M. R. (2016). The effects of time-varying temperature on delays in genetic networks. SIAM Journal on Applied Dynamical Systems, 15(3):1734–1752.

[Guo et al., 2014] Guo, S., Hori, Y., and Murray, R. (2014). Systematic design and implementation of a novel synthetic fold-change detector biocircuit in vivo. California Institute of Technology, Tech. Rep.

[Hart et al., 2014] Hart, Y., Reich-Zeliger, S., Antebi, Y. E., Zaretsky, I., Mayo, A. E., Alon, U., and Friedman, N. (2014). Paradoxical signaling by a secreted molecule leads to homeostasis of cell levels. Cell, 158(5):1022–1032.

[Hemme et al., 2010] Hemme, C. L., Mouttaki, H., Lee, Y.-J., Zhang, G., Goodwin, L., Lucas, S., Copeland, A., Lapidus, A., del Rio, T. G., Tice, H., et al (2010). Sequencing of multiple clostridial genomes related to biomass conversion and biofuel production. Journal of Bacteriology, 192(24):6494–6496.

[Hsiao et al., 2014] Hsiao, V., de los Santos, E. L., Whitaker, W. R., Dueber, J. E., and Murray, R. M. (2014). Design and implementation of a biomolecular concentration tracker. ACS synthetic biology, 4(2):150–161.

[Hsiao et al., 2016] Hsiao, V., Hori, Y., Rothemund, P. W., and Murray, R. M. (2016). A population-based temporal logic gate for timing and recording chemical events. Molecular Systems Biology, 12(5):869.

[Jishage and Ishihama, 1999] Jishage, M. and Ishihama, A. (1999). Transcriptional organization and in vivo role of the escherichia coli rsd gene, encoding the regulator of rna polymerase sigma d. Journal of Bacteriology, 181(12):3768–3776.

[Kampranis et al., 1999] Kampranis, S. C., Howells, A. J., and Maxwell, A. (1999). The interaction of dna gyrase with the bacterial toxin ccdb: Evidence for the existence of two gyrase-ccdb complexes. Journal of Molecular Biology, 293(3):733–744.

[Khalil and Collins, 2010] Khalil, A. S. and Collins, J. J. (2010). Synthetic biology: Applications come of age. Nature Reviews Genetics, 11(5):367.

[Klumpp et al., 2009] Klumpp, S., Zhang, Z., and Hwa, T. (2009). Growth rate-dependent global effects on gene expression in bacteria. Cell, 139(7):1366–1375.

[Kuhlman et al., 2003] Kuhlman, B., Dantas, G., Ireton, G. C., Varani, G., Stoddard, B. L., and Baker, D. (2003). Design of a novel globular protein fold with atomic-level accuracy. Science, 302(5649):1364–1368.

[Kwok, 2010] Kwok, R. (2010). Five hard truths for synthetic biology. Nature News, 463(7279):288–290.

[Levine, 2010] Levine, W. S. (2010). The Control Systems Handbook: Control System Advanced Methods. CRC press.

[Lewin, 2004] Lewin, B. (2004). Genes VIII. Number (Sirsi) i9780131439818. Pearson Prentice Hall Upper Saddle River, NJ.

[Lillacci et al., 2017] Lillacci, G., Aoki, S. K., Schweingruber, D., and Khammash, M. (2017). A synthetic integral feedback controller for robust tunable regulation in bacteria. bioRxiv, page 170951.

[Liu et al., 2012] Liu, C. C., Qi, L., Lucks, J. B., Segall-Shapiro, T. H., Wang, D., Mutalik, V. K., and Arkin, A. P. (2012). An adaptor from translational to transcriptional control enables predictable assembly of complex regulation. Nature methods, 9(11):1088–1094.

[Mathews et al., 2000] Mathews, C. K., Van Holde, K. E., and Ahern, K. G. (2000). Biochemistry. Add. Wesley Longman, San Francisco.

[McBride et al., 2003] McBride, M. T., Gammon, S., Pitesky, M., O’Brien, T. W., Smith, T., Aldrich, J., Langlois, R. G., Colston, B., and Venkateswaran, K. S. (2003). Multiplexed liquid arrays for simultaneous detection of simulants of biological warfare agents. Analytical Chemistry, 75(8):1924–1930.

[McCardell et al., 2017] McCardell, R. D., Huang, S., Green, L. N., and Murray, R. M. (2017). Control of bacterial population density with population feedback and molecular sequestration. bioRxiv, page 225045.

[Miller et al., 2011] Miller, C., Schwalb, B., Maier, K., Schulz, D., Dümcke, S., Zacher, B., Mayer, A., Sydow, J., Marcinowski, L., Dölken, L., et al (2011). Dynamic transcriptome analysis measures rates of mrna synthesis and decay in yeast. Molecular systems biology, 7(1):458.

[Moon et al., 2010] Moon, T. S., Dueber, J. E., Shiue, E., and Prather, K. L. J. (2010). Use of modular, synthetic scaffolds for improved production of glucaric acid in engineered e. coli. Metabolic engineering, 12(3):298–305.

[Moon et al., 2012] Moon, T. S., Lou, C., Tamsir, A., Stanton, B. C., and Voigt, C. A. (2012). Genetic programs constructed from layered logic gates in single cells. Nature, 491(7423):249.

[Narcross et al., 2016] Narcross, L., Fossati, E., Bourgeois, L., Dueber, J. E., and Martin, V. J. (2016). Microbial factories for the production of benzylisoquinoline alkaloids. Trends in Biotechnology, 34(3):228–241.

[Olsman et al., 2018] Olsman, N., Baetica, A.-A., Xiao, F., Leong, Y. P., Doyle, J., and Murray, R. (2018). Hard limits and performance tradeoffs in a class of sequestration feedback systems. bioRxiv.

[Pardee et al., 2014] Pardee, K., Green, A. A., Ferrante, T., Cameron, D. E., DaleyKeyser, A., Yin, P., and Collins, J. J. (2014). Paper-based synthetic gene networks. Cell, 159(4):940–954.

[Purnick and Weiss, 2009] Purnick, P. E. and Weiss, R. (2009). The second wave of synthetic biology: From modules to systems. Nature Reviews Molecular Cell Biology, 10(6):410.

[Qian et al., 2011] Qian, L., Winfree, E., and Bruck, J. (2011). Neural network computation with dna strand displacement cascades. Nature, 475(7356):368.

[Qian and Del Vecchio, 2018] Qian, Y. and Del Vecchio, D. (2018). Realizing ‘integral control’ in living cells: How to overcome leaky integration due to dilution? Journal of The Royal Society Interface, 15(139).

[Rajasekar et al., 2016] Rajasekar, K. V., Zdanowski, K., Yan, J., Hopper, J. T., Francis, M.-L. R., Seepersad, C., Sharp, C., Pecqueur, L., Werner, J. M., Robinson, C. V., et al (2016). The anti-sigma factor rsra responds to oxidative stress by reburying its hydrophobic core. Nature Communications, 7:12194.

[Ren et al., 2017] Ren, X., Baetica, A.-A., Swaminathan, A., and Murray, R. M. (2017). Population regulation in microbial consortia using dual feedback control. In Decision and Control (CDC), 2017 IEEE 56th Annual Conference on, pages 5341–5347. IEEE.

[Rust et al., 2007] Rust, M. J., Markson, J. S., Lane, W. S., Fisher, D. S., and O’Shea, E. K. (2007). Ordered phosphorylation governs oscillation of a three-protein circadian clock. Science, 318(5851):809–812.

[Savage et al., 2008] Savage, D. F., Way, J., and Silver, P. A. (2008). Defossiling fuel: How synthetic biology can transform biofuel production. ACS Chem. Biol.

[Sharma and Chatterji, 2008] Sharma, U. K. and Chatterji, D. (2008). Differential mechanisms of binding of anti-sigma factors escherichia coli rsd and bacteriophage t4 asia to e. coli rna polymerase lead to diverse physiological consequences. Journal of bacteriology, 190(10):3434–3443.

[Steif and Meyer, 2012] Steif, A. and Meyer, I. M. (2012). The hok mrna family. RNA Biology, 9(12):1399–1404.

[Thubagere Jagadeesh, 2017] Thubagere Jagadeesh, A. (2017). Programming Complex Behavior In DNA-based Molecular Circuits And Robots. PhD thesis, California Institute of Technology.

[Walton et al., 2002] Walton, S. P., Stephanopoulos, G. N., Yarmush, M. L., and Roth, C. M. (2002). Thermodynamic and kinetic characterization of antisense oligodeoxynucleotide binding to a structured mrna. Biophysical journal, 82(1):366–377.

[Wang et al., 2009] Wang, L., Chen, W., Xu, D., Shim, B. S., Zhu, Y., Sun, F., Liu, L., Peng, C., Jin, Z., Xu, C., et al (2009). Simple, rapid, sensitive, and versatile swnt-paper sensor for environmental toxin detection competitive with elisa. Nano letters, 9(12):4147–4152.

[Werner, 2010] Werner, J. (2010). System properties, feedback control and effector coordination of human temperature regulation. European Journal of Applied Physiology, 109(1):13–25.

[Xiao and Doyle, 2018] Xiao, F. and Doyle, J. C. (2018). Robust perfect adaptation in biomolecular reaction networks. bioRxiv.

[Yeung et al., 2017] Yeung, E., Dy, A. J., Martin, K. B., Ng, A. H., Del Vecchio, D., Beck, J. L., Collins, J. J., and Murray, R. M. (2017). Biophysical constraints arising from compositional context in synthetic gene networks. Cell Systems, 5(1):11–24.

[Yeung et al., 2014] Yeung, E., Ng, A., Kim, J., Sun, Z. Z., and Murray, R. M. (2014). Modeling the effects of compositional context on promoter activity in an e. coli extract based transcription-translation system. In Decision and Control (CDC), 2014 IEEE 53rd Annual Conference on, pages 5405–5412. IEEE.

[Yi et al., 2000] Yi, T.-M., Huang, Y., Simon, M. I., and Doyle, J. (2000). Robust perfect adaptation in bacterial chemotaxis through integral feedback control. Proceedings of the National Academy of Sciences, 97(9):4649–4653.

[Zhou and Gottesman, 1998] Zhou, Y. and Gottesman, S. (1998). Regulation of proteolysis of the stationary-phase sigma factor rpos. Journal of Bacteriology, 180(5):1154–1158.

